# Clonally-Expanded, Thyrotoxic Autoimmune Mediator CD8^+^ T cells Driven by IL21 Contribute to Checkpoint Inhibitor Thyroiditis

**DOI:** 10.1101/2022.12.18.517398

**Authors:** Melissa G. Lechner, Zikang Zhou, Aline T. Hoang, Nicole Huang, Jessica Ortega, Lauren N. Scott, Ho-Chung Chen, Anushi Y. Patel, Rana Yakhshi-Tafti, Kristy Kim, Willy Hugo, Pouyan Famini, Alexandra Drakaki, Antoni Ribas, Trevor E. Angell, Maureen A. Su

## Abstract

Autoimmune toxicity occurs in up to 60% of patients treated with immune checkpoint inhibitor (ICI) cancer therapy and is an increasing clinical challenge with the expanding use of these treatments. To date, human immunopathogenic studies of immune related adverse events (IRAEs) have relied upon sampling of circulating peripheral blood cells rather than affected tissues. Here, we directly obtained thyroid specimens from subjects with ICI-thyroiditis, one of the most common IRAEs, and compared immune infiltrates to those from subjects with spontaneous autoimmune Hashimoto’s thyroiditis (HT) or no thyroid disease. Single cell RNA sequencing revealed a dominant, clonally expanded population of thyroid-infiltrating cytotoxic CXCR6^+^ CD8^+^ T cells (“CD8^+^ autoimmune mediators) present in ICI-thyroiditis, but not HT or healthy controls. Furthermore, we identified a crucial role for interleukin 21, a cytokine secreted by intrathyroidal T follicular (Tfh) and T peripheral helper (Tph) cells, as a driver of these thyrotoxic CD8^+^ autoimmune mediators. In the presence of IL21, human CD8^+^ T cells acquired the autoimmune mediator phenotype with upregulation of cytotoxic molecules (IFNγ, granzyme); the chemokine receptor CXCR6; and thyrotoxic capacity. We validated these findings *in vivo* using a novel mouse model of IRAEs, and further demonstrated that genetic blockade of IL21 signaling protected ICI-treated mice from thyroid immune infiltration. Taken together these studies reveal novel mechanisms and therapeutic targets by which IL21^+^ Tfh/Tph cells drive thyrotoxic CD8^+^ autoimmune mediators for the development of IRAEs in humans.

**One Sentence Summary:** ScRNAseq reveals a novel role for CD8^+^ autoimmune mediators and IL21^+^ T helper cells in the pathogenesis of human checkpoint inhibitor thyroiditis.

## INTRODUCTION

Immune checkpoint inhibitors (ICI) [*e.g*. anti-programmed death protein/ligand (PD1/PDL1), anti-cytotoxic T lymphocyte antigen (CTLA4)] have dramatically improved outcomes for many patients with advanced malignancies *(1)*. But with increased immune activation can come unwanted autoimmune attack on healthy tissues. Such immune related adverse events (IRAEs) occur in up to 60% of patients treated with ICI cancer therapy and can contribute to treatment interruption, hospitalizations, and even premature death *(2, 3)*. With the expanding use of ICI therapy *(4)*, IRAEs are an increasing clinical problem.

The thyroid is one of the most common organs affected by ICI-associated autoimmunity, targeted in approximately 10-15% of patients treated with anti-PD1/L1 monotherapy and nearly 30% of patients treated with combination anti-PD1/L1 and CTLA4 *(5–7)*. ICI-induced thyroid autoimmunity, or thyroiditis, presents as immune infiltration and rapid destruction of the thyroid gland, leading to hormone abnormalities and requiring lifelong thyroid hormone replacement in most patients *(5, 6, 8)*. Despite significant efforts to date, the cause of many IRAEs remain poorly understood, including ICI-thyroiditis. Recent mechanistic studies by us and others showed a key role for T cells and interleukin 17A (IL17A) in ICI-thyroiditis in mouse models *(9–11)*. It remains unclear, however, whether these findings translate to human IRAEs, which highlights the need for evaluating IRAE immunopathogenesis in human subjects.

Like IRAEs in other organs, ICI-thyroiditis has both overlapping and distinct features with spontaneous autoimmunity (*i.e*. Hashimoto’s thyroiditis, HT) *(6–8, 12)*. HT is a common form of thyroid autoimmunity present in nearly 10% of the US population. HT is characterized by slowly progressive thyroid gland immune cell-mediated destruction *(13)*. Central to the pathogenesis of HT are Type 3 immune responses, namely IL17 producing CD4^+^ T helper cells *(14–17)*. In addition, tertiary lymphoid follicular structures are classically seen in thyroid immune infiltrates in HT *(18)*, in conjunction with increased circulating T follicular helper cells compared to healthy controls *(19)*. Two prior studies *(20, 21)* showed intrathyroidal lymphocyte accumulation in the thyroid tissue of patients with ICI-thyroiditis, similar to HT.

On the other hand, ICI-thyroiditis has several notable differences from HT. In contrast to spontaneous HT, thyroid gland destruction in ICI-thyroiditis is more rapid, occurring over weeks rather than years *(6, 8)*. Furthermore, approximately half of patients with ICI-thyroiditis lack the hallmark anti-thyroid antibodies present in HT *(6, 8)*. Thus, ICI-thyroiditis, while likely sharing some overlapping mechanisms with HT, may also be driven by additional distinct immune mechanisms.

To develop strategies to reduce ICI autoimmune toxicities in patients, driving mechanisms need to be identified. Studies into the immunopathogenesis of IRAEs in humans have largely relied upon sampling of peripheral blood rather than affected tissues *(22–25)*. Therefore, to delineate human IRAE pathogenesis, we sampled thyroid tissue from subjects with ICI-thyroiditis and compared them to those from individuals with HT and healthy controls. Using single cell RNA sequencing of human thyroid fine needle aspiration (FNA) specimens, we identify CXCR6^+^ interferon gamma (IFNγ)^+^ cytotoxic CD8^+^ T cells [“CD8^+^ autoimmune mediators” *(26–28)*] as key contributors to ICI-thyroiditis (IRAE). In addition, we identify a mechanism by which interleukin 21, an important cytokine secreted by T follicular (Tfh) and T peripheral helper (Tph) cells, can promote differentiation of these thyrotoxic CD8^+^ autoimmune mediators. This role of IL21^+^ Tfh and Tph cells, which were present in both HT and ICI-thyroiditis, suggests a potential link between underlying thyroid autoimmunity and the development of thyroid IRAEs. Finally, we corroborate these findings *in vivo* using a novel mouse model of ICI-associated autoimmunity and show that genetic blockade of IL21 receptor signaling reduced autoimmune mediator numbers and thyroid autoimmune infiltrates. Together, these findings demonstrate a key thyrotoxic pathway in which IL21^+^ Tfh/Tph cells drive CD8^+^ autoimmune mediator development in IRAEs.

## RESULTS

### Thyroid-infiltrating immune populations in ICI-thyroiditis patients are T cell-predominant and include IL21^+^ CD4^+^ Tfh and Tph cells

T cells are presumed mediators of ICI-related autoimmunity. However, these conclusions are based largely on data derived from circulating immune populations, inferences from anti-tumor immune responses, or preclinical models *(1, 9, 10, 22–24, 29, 30)*. Direct data from affected tissues of patients with IRAEs have been limited *(20, 21)*. Therefore, to better understand the mechanisms driving IRAEs in humans, we evaluated immune cells in thyroid specimens from patients with ICI-thyroiditis (IRAE, n=9 patients) (**Fig. 1A**). IRAE patients included recipients of combination anti-PD1 and anti-CTLA4, anti-PD1, or anti-PD-L1 therapies for non-thyroid, solid malignancies and developed newly abnormal thyroid function tests consistent with thyroiditis (suppressed TSH and elevated FT4). Thyroid fine needle aspirates were collected within 2 months of the first abnormal thyroid function tests and thyroid autoantibodies (anti-thyroglobulin or anti-thyroid peroxidase) were present in 4/9 patients. Full clinical and demographic parameters for patients are shown in **Table S1**.

**Fig. 1.**
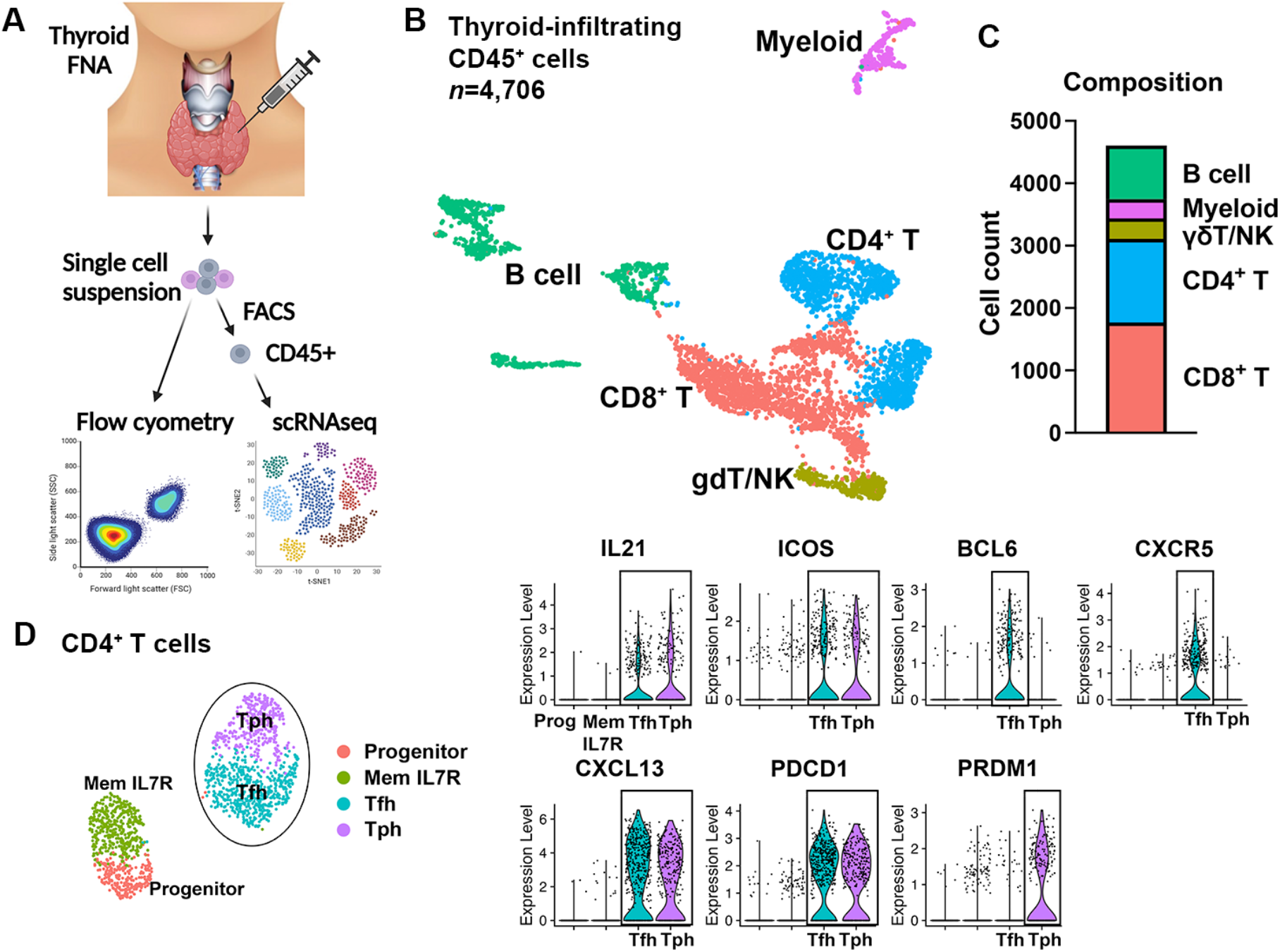
Thyroid-infiltrating immune populations in ICI-thyroiditis patients are T cell-predominant and include IL21^+^ CD4^+^ Tfh and Tph cells. **(A)** Schema of thyroid specimen collection by fine needle aspiration (FNA). **(B)** Single cell RNA sequencing analysis of thyroid infiltrating CD45^+^ immune cells (*n*=5 patients), UMAP shown. **(C)** Distribution of intrathyroidal cells by major immune population. **(D)** UMAP of subclustered CD4^+^ T cells (*left*) and violin plots showing genes associated with T follicular (Tfh) and peripheral (Tph) helper populations (*right*).

Given the small number of cells present in thyroid specimens (approximately 10,000 cells per aspirate), we first sought to ascertain that immune cells could be detected within these samples. Flow cytometry analysis showed that immune cells were indeed present, and that T cells were the primary immune cell subset in ICI-thyroiditis (**Fig. S1A**), consistent with a prior report by Kotwal *et al*. evaluating thyroid IRAEs *(20)*. We then turned to single cell RNA sequencing (scRNAseq) for a more detailed evaluation of immune infiltrates (**Fig. 1B**). CD45^+^ thyroid infiltrating immune cells were sorted from thyroid specimens (n=5 IRAE patients) and subjected to the 10x Genomics scRNAseq pipeline. Uniform Manifold Approximation and Projection (UMAP) visualization revealed 14 distinct groups (**Fig. S1B and Data set S1**), representing six broad cell types: CD4^+^ T cells (*CD3E, CD4*), CD8^+^ T cells (CD8: *CD3, CD8A, CD8B*), gamma delta (γδ) T cells (*CD3E, TRDC*, variable *TRGV* genes), B cells (*CD19, CD79A*), myeloid (*CD14, CD68, CSF1R*) and natural killer (NK) cells (*NKG7, NCR1, FGFBP2*) (**Fig. 1B**). By absolute cell count, CD4^+^ and CD8^+^ T cells make up the majority of immune cells infiltrating the thyroid (**Fig. 1C**). These data demonstrate that thyroid immune infiltrates in ICI-thyroiditis patients contain diverse immune cells, with T cells as the primary population.

CD4^+^ T helper cells drive spontaneous thyroid autoimmunity in many tissues *(15, 16, 19)* and have been implicated in ICI-associated thyroid autoimmunity in mouse models *(9, 10, 29)*. Therefore, we took a closer look at thyroid-infiltrating CD4^+^ T cells in our scRNAseq data (**Fig. 1D**). Interestingly, thyroid infiltrates were composed largely of T follicular (Tfh, *PDCD1*^*+*^ *ICOS*^*+*^ *BCL6*^*+*^ *CXCR5*^*+*^) and T peripheral helper cells (Tph, *PDCD1*^*+*^ *ICOS*^*+*^ *CXCR5*^*-*^ *PRDM1*^*+*^), populations that have been associated with multiple spontaneous autoimmune diseases, including HT *(19, 31–34)*. Tfh and Tph classically recruit B and T cells via chemokine CXCL13 and produce IL21 to promote B cell antibody production *(32, 33)* (**Fig. 1D**). In addition, recent data suggested that IL21 from Tfh may drive increased pathogenicity of CD8^+^ T cells in anti-viral and anti-tumor immune responses *(35–37)*.

### Comparison to Hashimoto’s thyroiditis reveals expansion of CD8^+^ T cells in ICI-thyroiditis

Based upon the presence of CD4^+^ Tfh and Tph in ICI-thyroiditis, populations previously associated with spontaneous autoimmunity, we sought to probe the relationship between ICI-thyroiditis and HT. Using scRNAseq, we compared thyroid immune infiltrates in ICI-thyroiditis patients to those with HT (n=5), as well as controls with no thyroid disease (Healthy Control; n=3). HT patients were defined as having thyroid autoantibodies, ultrasound imaging changes consistent with chronic lymphocytic thyroiditis, and biochemical hypothyroidism and/or treatment with levothyroxine (**Table S1**). Healthy controls were biochemically and clinically euthyroid, had no thyroid autoantibodies, and normal thyroid ultrasound appearance.

We integrated scRNAseq data from IRAE, HT, and healthy control thyroid specimens and visualized the data by UMAP. Clustering of CD45^+^ intrathyroidal immune cells again showed diverse immune populations, with 13 distinct clusters that were represented across all conditions (**Fig. 2A**; differentially expressed genes in **Data set S1 and Fig. S2A**). Similar to ICI-thyroiditis alone, CD4^+^ (*CD3E, CD4*) and CD8^+^ T cells (*CD3E, CD8A, CD8B)* dominated immune infiltrates (**Fig. S2A**). Other populations were B (CD19, CD79A), myeloid (*CD14, CD68*), gamma delta (γδ) T cells (*CD3E, TRDC*, variable *TRGV* genes), and NK cells (*NKG7, FCGR3A, NCR1*) (**Fig. S2B**). Importantly, *IL21*^+^ CD4^+^ T cells (cluster CD4-4), comprising Tfh and Tph cells, were prominent in thyroid immune infiltrates and present across both HT and ICI-thyroiditis states (**Fig. 2A and Fig. S2A)**. These data are consistent with a prior report by Zhu et al. *(19)* showing increased Tfh in HT, and support a role for IL21^+^ CD4^+^ cells in ICI-thyroiditis. Thus, IL21^+^ CD4^+^ T cells may be a shared mechanism between HT and ICI-thyroiditis.

**Fig. 2.**
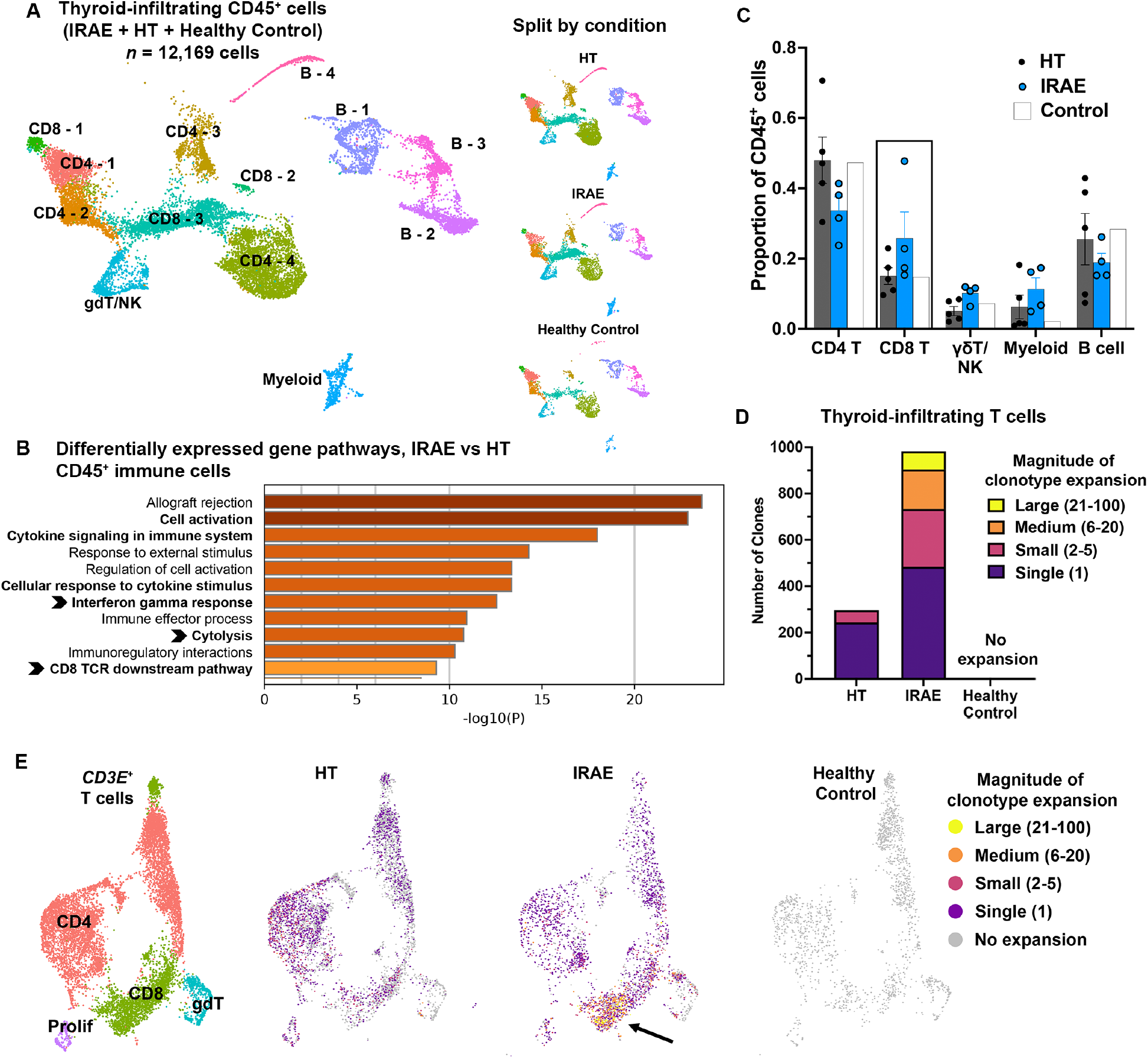
CD8^+^ T cells are expanded in ICI-thyroiditis compared to Hashimoto’s. **(A)** Comparison of thyroid immune infiltrates in ICI-thyroiditis (IRAE), Hashimoto’s thyroiditis (HT) and healthy controls by single cell RNA sequencing. UMAP of integrated CD45^+^ cells across conditions (*left*), with UMAP plots split by condition on *right*. **(B)** Pathway analysis of differentially expressed genes in thyroid-infiltrating CD45^+^ cells between IRAE and HT. **(C)** Distribution of intrathyroidal cells by major immune population by condition. ANOVA, p=ns **(D)** Frequency of clonotype expansion of intrathyroidal CD3^+^ T cells by thyroid state. **(E)** UMAP of thyroid-infiltrating CD3^+^ cells (*left*) and clonotype distribution split by thyroid condition (*right*).

To identify immune mechanisms unique to IRAEs, we performed differential pathway analysis between HT and ICI-thyroiditis specimens. CD8^+^ T cell-related pathways, notably CD8 T cell receptor (TCR) signaling, cytolysis, and interferon gamma (IFNγ) response, were significantly increased in ICI-thyroiditis compared to HT (**Fig. 2B**). Consistent with this, CD8^+^ T cells were also increased in frequency among thyroid-infiltrating immune cells in ICI-thyroiditis compared to HT (**Fig. 2C**).

The expansion of specific T cell receptor clonotypes is an indicator of antigen recognition and subsequent activation and proliferation among CD8^+^ T cells. We queried T cell clonal expansion in thyroid immune infiltrates across thyroid states using 10x single cell TCR sequencing of thyroid specimens from IRAE (n=5), HT (n=5), and healthy control (n=3) subjects. Overall T cell clonal expansion was greater in patients with ICI-thyroiditis compared to HT, and no clonal expansion was seen in healthy controls (**Fig. 2D**). Clonally expanded T cells mapped to multiple CD4^+^ and CD8^+^ populations, but the largest clonotype expansion in ICI-thyroiditis was seen in CD8^+^ T cells (**Fig. 2E**). Together, these data suggest a more prominent role of clonally-expanded CD8^+^ T cells in the pathogenesis of ICI-thyroiditis.

### ICI-thyroiditis is distinguished from HT by intrathyroidal accumulation of clonally expanded, CXCR6^+^ GZMB^+^ IFNG^+^ CD8^+^ “autoimmune mediator” T cells

To further explore how CD8^+^ T cells may be contributing to IRAEs, we subsetted out CD8^+^ T cells from the larger population of thyroid infiltrating CD45^+^ immune cells and compared gene expression across thyroid conditions. Interestingly, CD8^+^ T cells in patients with ICI-thyroiditis showed significantly greater expression of *IFNG, GZMB, FASLG*, and *CXCR6* than HT or healthy controls (**Fig. 3A**). These genes drive cell-mediated immunity, cytotoxicity, and T cell homing to inflamed tissues *(38–42)* – key processes in autoimmune tissue attack. By contrast, genes associated with a progenitor phenotype, including *TCF7* (encoding TCF1) and *SELL*, had lower expression in ICI-thyroiditis (**Fig. 3A)**. These data suggest that effector CD8^+^ T cells are a critical component of ICI-thyroiditis and distinguish this disease entity from other thyroid states.

**Fig. 3.**
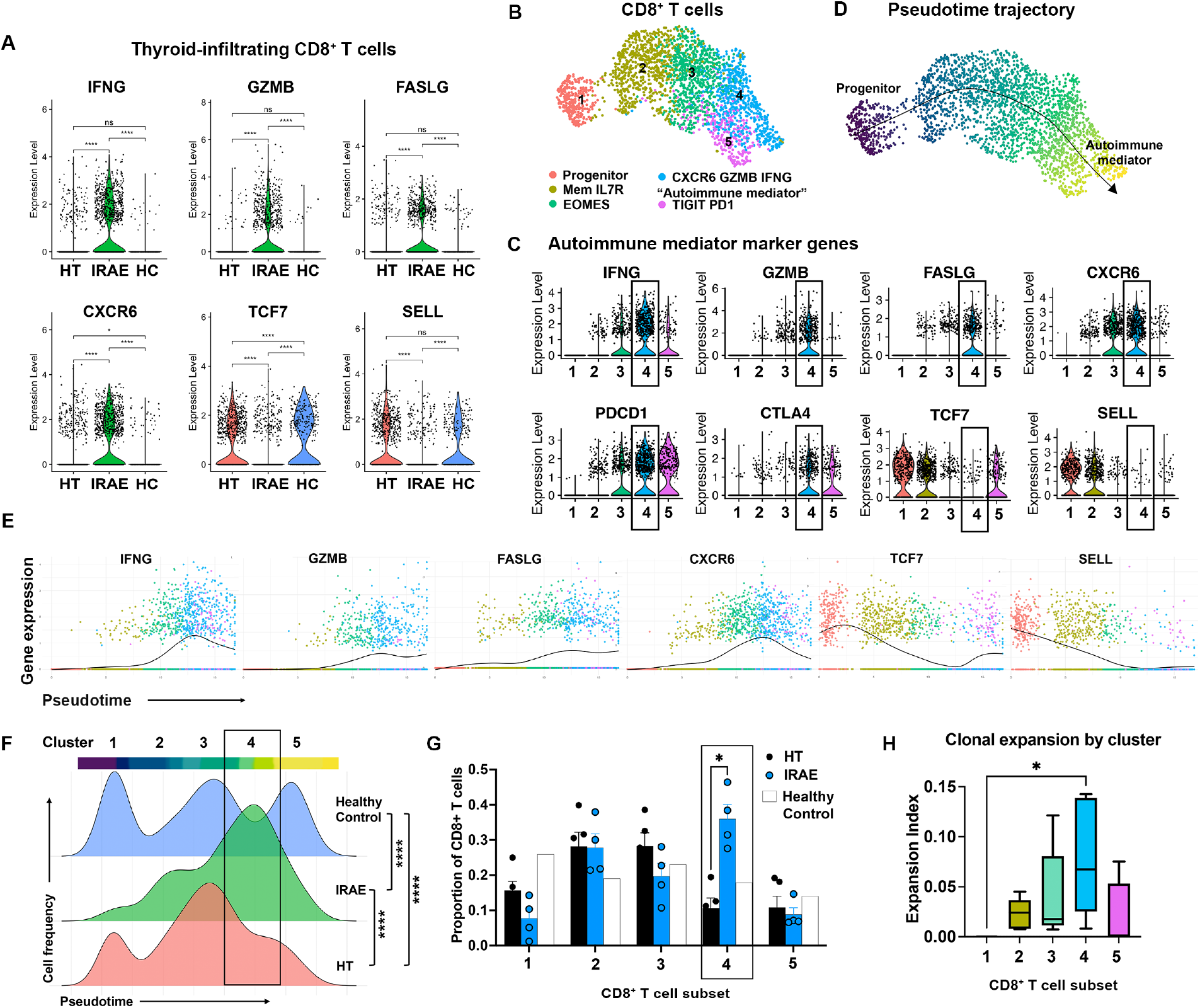
Clonally expanded CD8^+^ autoimmune mediator T cells distinguish ICI-thyroiditis. **(A)** CD8^+^ T cell gene expression by thyroid state. Hashimoto’s (HT), ICI-thyroiditis (IRAE), and healthy controls (HC). **(B)** UMAP of subclustered thyroid-infiltrating CD8^+^ T cells. **(C)** Gene expression across CD8^+^ subclusters. **(D)** Slingshot trajectory analysis of CD8^+^ T cells. **(E)** Transcriptional transitions in CD8^+^ T cells along the pseudotime trajectory. **(F)** Density distribution of CD8^+^ T cell along the psuedotime trajectory, split by thyroid state. **(G)** Comparison of subset frequency between IRAE and HT; pooled HC specimen also shown. **(H)** STARTRAC clonal expansion index by CD8^+^ subcluster. ANOVA with Welch correction (A, F), asymptotic Kolmogorov-Smirnov test (F), or Student’s t test with correction for multiple comparisons (G), *p<0.05, ***p<0.001, ****p<0.0001.

We subclustered CD8^+^ T cells to better define the specific populations comprising autoimmune infiltrates in the thyroid. Intrathyroidal CD8^+^ T cells were comprised of five distinct groups (**Fig. 3B**, and **Data set S1**), defined as progenitor (Cluster 1; *SELL, TCF7*), memory IL7R (Cluster 2; *IL7R, SELL, TCF7*), EOMES (Cluster 3; *EOMES*), CXCR6 (Cluster 4; *CXCR6, IFNG, GZMB*) or TIGIT (Cluster 5; *TIGIT, LAG3*). Notably, subclustering revealed a population of CD8^+^ T cells in Cluster 4 with a gene signature that mirrored the differentially expressed genes in CD8^+^ T cells from ICI-thyroiditis vs HT and HC, namely *CXCR6*^+^*GZMB*^+^ *FASLG*^+^ *PDCD1*^+^ and *TCF7*^-^ (**Fig. 3C**). Furthermore, the cells in Cluster 4 matched the phenotype of “CD8^+^ autoimmune mediators” that have recently been associated with autoimmunity. Gearty et al. *(28)* and Ciecko et al. *(27)* both identified IFNG^+^ GZM^+^ CXCR6^+^ TCF1^-^ CD8^+^ T cells as immune effectors contributing to autoimmune tissue attack in the pancreas in mouse models of diabetes mellitus. In addition, Dudek and colleagues reported that cytotoxic CXCR6^+^ CD8^+^ autoimmune mediator T cells can drive immune-mediated liver injury *(26)*. However, CD8^+^ autoimmune mediator T cells have not previously been associated with autoimmune thyroiditis in humans.

Whether CD8^+^ autoimmune mediators migrate from draining lymph nodes or develop within inflamed tissues is unclear *(27, 28)*. To define the ontogeny of these CD8^+^ autoimmune mediator T cells, we examined transcriptional transitions among intrathyroidal CD8^+^ subsets. Trajectory analysis showed that CD8^+^ T cells progressed along a single lineage from progenitors to autoimmune mediators (**Fig. 3D**). Specifically, CD8^+^ cells decreased expression of *TCF7* and *SELL* and increased expression of *IFNG, GZMB, FASLG*, and *CXCR6* along the trajectory (**Fig. 3E**). This is most consistent with the findings from Ceicko et al. *(27)* in diabetes showing that progenitor *CXCR6*^-^ *TCF7*^+^ CD8^+^ T cells give rise to pathogenic *CXCR6*^+^ *TCF7*^-^ CD8^+^ autoimmune mediator T cells in the tissue and that these progeny are sufficient to cause autoimmunity. Thus, CD8^+^ autoimmune mediators in human thyroiditis arise from intrathyroidal progenitors via a differentiation pathway that may be shared across autoimmune diseases.

Importantly, intrathyroidal CD8^+^ T cell differentiation was distinct across thyroid states. In ICI-thyroiditis patients, the majority of CD8^+^ T cells showed differentiation to mature CD8^+^ autoimmune mediators (CXCR6^+^ GZMB^+^ IFNG^+^TCF7^-^) (**Fig. 3F**). In contrast, CD8^+^ T cells in healthy controls remained primarily in the progenitor or exhausted TIGIT^+^ PD1^+^ subsets, while CD8^+^ T cells in HT patients were most likely in the EOMES^+^ cluster as precursors to the autoimmune mediator population. Consistent with these data, ICI-thyroiditis patients had a greater frequency of CD8^+^ autoimmune mediators (Cluster 4) in thyroid immune infiltrates than HT patients (**Fig. 3G**, p<0.05).

Notably, TCR clonal expansion, an indicator of antigen recognition and response within CD8^+^ T cell populations, was greatest within the CD8^+^ autoimmune mediators (Cluster 4) (**Fig. 3H**), further supporting their participation in thyroid tissue autoimmune attack. In addition, greater clonotype expansion correlated with increasing expression of cytotoxic and effector genes (*GZMB, FASLG, IFNG, CXCR6*) (**Fig. S3A)**. Shared clonotypes across clusters also serve as lineage markers as cells undergo transcriptional transitions. Clonally expanded CD8^+^ autoimmune mediator cells (Cluster 4) had shared clonotypes with cells across the EOMES^+^ (Cluster 3) and TIGIT^+^ (Cluster 5) subsets (**Fig. S3B)**. In addition, the clonal transition index was highest across these three clusters (**Fig. S3C**). Taken together, this supports the notion that autoimmune mediators may be recruited from other subsets in the thyroid during ICI therapy. Thus, our data show a direct relationship between clonal expansion and acquisition of cytotoxic effector function by CD8^+^ autoimmune mediator T cells in thyroid immune infiltrates. In summary, our data show that a distinguishing feature of ICI-thyroiditis is the differentiation of clonally-expanded CD8^+^ T cells to cytotoxic autoimmune mediators in the thyroid.

### IL21 promotes the CD8^+^ T cell autoimmune mediator phenotype

Our initial observations in IRAE samples showed prominent *IL21*^*+*^ CD4^+^ Tfh and Tph cells in thyroid immune infiltrates (**Fig. 1D**). How CD4^+^ Tfh and Tph cell population dynamics change between IRAE, HT, and healthy controls, however, remain unclear. Using our integrated data set of thyroid-infiltrating immune cells from IRAE, HT, and healthy control subjects, we isolated and subclustered the CD4^+^ T cells. We delineated thyroid-infiltrating CD4^+^ T cell subsets across thyroid conditions, yielding eight distinct populations (**Fig. 4A**). In addition to three Tfh and Tph clusters [Tfh-1 (*ICOS, CXCL13, PDCD1*), Tfh-2 (*TOX2, ICOS, CXCL13, PDCD1, CXCR5, IL21*) and Tph (*ICOS, PDCD1*)], we also saw CCR7 (*SELL, CCR7, LEF1*), memory IL7R (*IL7R, SELL*), IL7R PD1 (*IL7R, PDCD1, CD44*), CD44, and GZMK groups (**Data set S1**). Importantly, Tfh and Tph cells expressed high levels of *IL21, CXCL13, PDCD1* and *CTLA4* (**Fig. 4B**).

**Fig. 4.**
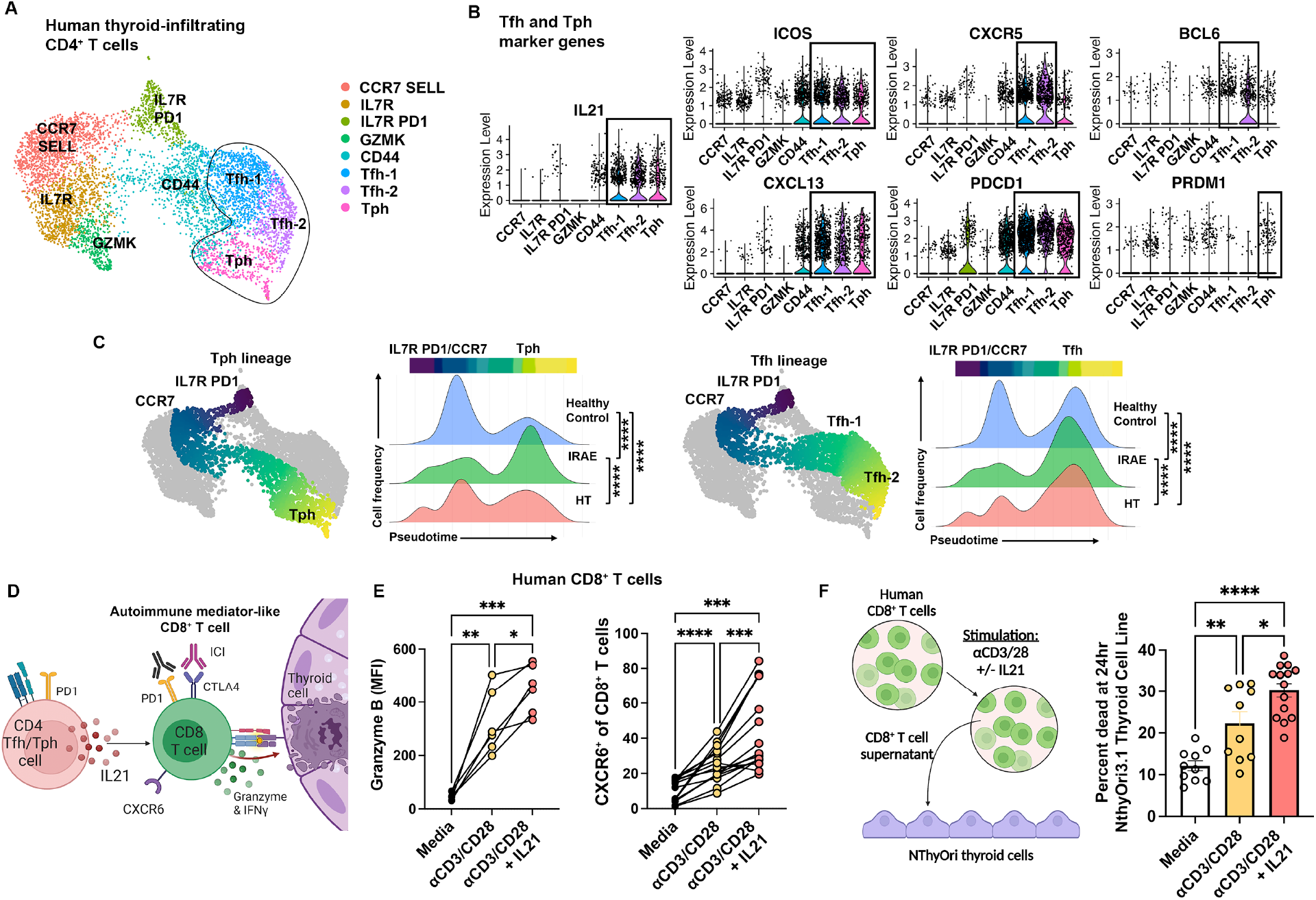
IL21^+^ CD4^+^ T follicular helper (Tfh) and peripheral helper (Tph) cells are enriched in ICI-thyroiditis and promote CD8^+^ T cell thyrotoxicity. **(A)** UMAP of subclustered thyroid infiltrating CD4^+^ T cells in human FNA specimens by scRNAseq. **(B)** Expression of Tfh and Tph marker genes by cluster. **(C)** Trajectory analysis of CD4^+^ T cells by Slingshot and density distribution along the psuedotime trajectory by condition for two lineages: Tph (*left*) and Tfh (*right*). **(D)** Proposed action of Tfh and Tph cells on CD8^+^ autoimmune mediators in ICI-thyroiditis. **(E)** Quantification of granzyme B and CXCR6 expression on human CD8^+^ T cells following *in vitro* stimulation with anti-CD3/28 beads and recombinant human IL21, evaluated after 3 days by flow cytometry. **(F)** NThyOri3.1 human thyroid epithelial cell killing induced by incubation with supernatants from human CD8^+^ T cell cultures stimulated as indicated. Asymptotic Kolmogorov-Smirnov test (C), repeated measure ANOVA (E) or ANOVA with Welch correction (F), and subsequent pairwise comparisons. *p<0.05, **p<0.01, ***p<0.001, ****p<0.0001.

Furthermore, trajectory analysis showed that differentiation from IL7R PD1 and CCR7 cells to Tph was significantly increased in ICI-thyroiditis compared to HT and healthy controls (**Fig. 4C**, *left*). Differentiation from IL7R PD1 and CCR7 cells to Tfh cells was increased in both ICI-thyroiditis and HT compared to healthy controls (**Fig. 4C**). Thus, differentiation toward *IL21*^+^ CD4^+^ Tph and Tfh cells was increased within the thyroid tissue in ICI-thyroiditis patients, indicating their potential importance to IRAE immunopathogenesis.

Recent data show that, aside from its established role in promoting B cell responses, IL21 can promote CD8^+^ T cell pathogenicity *(35–37)*. These data led us to hypothesize that IL21 produced by Tfh and Tph may be promoting CD8^+^ autoimmune mediator T cell differentiation in ICI-thyroiditis (**Fig. 4D**. Indeed, *in vitro* stimulation with anti-CD3/28 and recombinant IL21 increased expression of the cytotoxic molecule granzyme B and the chemokine receptor CXCR6 in human CD8^+^ T cells, compared to anti-CD3/28 alone (**Fig. 4E**). IL21-induced CXCR6^+^ CD8^+^ T cells also expressed checkpoint proteins PD1 and CTLA4 (**Fig. S4A**), similar to autoimmune mediators described by Cieko et al. *(27)* and Dudek et al. *(26)*. Thus, IL21 promotes differentiation of human CD8^+^ T cells to become autoimmune mediator-like cells with a CXCR6^+^ GZM^+^ IFNG^+^ PD1^+^ CTLA4^+^ phenotype.

Despite their expression of cytotoxic molecules and effector molecules, it remains unclear whether IL21-induced CD8^+^ T cells are thyrotoxic. For these studies, we incubated human thyroid cells (NThyOri3.1 cell line) with supernatants from IL21-stimulated human CD8^+^ T cells. Importantly, incubation with supernatants from IL21 + anti-CD3/28-stimulated CD8^+^ T cells resulted in greater killing of human thyroid cells compared to supernatants from anti-CD3/28-treated or unstimulated CD8^+^ T cells (**Fig. 4F**). Thus, IL21 from Tfh and Tph cells can drive human CD8^+^ T cells to become thyrotoxic autoimmune mediators.

### ICI treatment in an IRAE mouse model recapitulates expansion of CD8^+^ autoimmune mediators and CD4^+^ Tfh/Tph cells

Our data from human thyroid specimens revealed a mechanism by which IL21 from Tfh and Tph cells contributes to the development of ICI-thyroiditis through the expansion of thyrotoxic CD8^+^ autoimmune mediators. To further explore this mechanism *in vivo*, we turned to our recently developed IRAE mouse model *(9)*. In this model, autoimmunity-prone non-obese diabetic (NOD) mice treated with anti-PD1 and/or anti-CTLA4 monoclonal antibodies develop multi-organ autoimmunity, including thyroiditis. Similar to patients treated with ICI for cancer therapy, mice developed autoimmunity more frequently with the combination of anti-PD1 + anti-CTLA4 (Dual ICI) *vs*. with single agent ICI treatment. Furthermore, NOD mice engrafted with MC38.β2M^-/-^ colon carcinoma or B16.β2M^-/-^ melanoma could be used to study ICI-associated autoimmunity in the context of anti-tumor immune responses (**Fig. 5A**). In our model, CD8^+^ autoimmune mediator (CXCR6^+^ GZMB^+^ IFNγ^+^) T cells were increased in ICI-treated NOD mice *vs*. isotype controls in both tumor models evaluated (**Fig. 5B**). Additionally, ICI-treated mice had increased Tfh (CXCR5^-^ PD1^+^ ICOS^+^ CD4^+^) and Tph (CXCR5^-^ PD1^+^ ICOS^+^ CD4^+^) cells (**Fig. 5C**) and greater IL21 production by helper T cells (**Fig. 5D**). These findings recapitulate the human finding that ICI treatment is associated with expansion of CD8^+^ autoimmune mediators, CD4^+^ Tfh cells, and CD4^+^ Tph populations. In summary, our data show that in patients with cancer and in an *in vivo* tumor-bearing mouse model there is an increase in IL21^+^ Tfh and Tph cells and CD8^+^ autoimmune mediator T cells during ICI therapy.

**Fig. 5.**
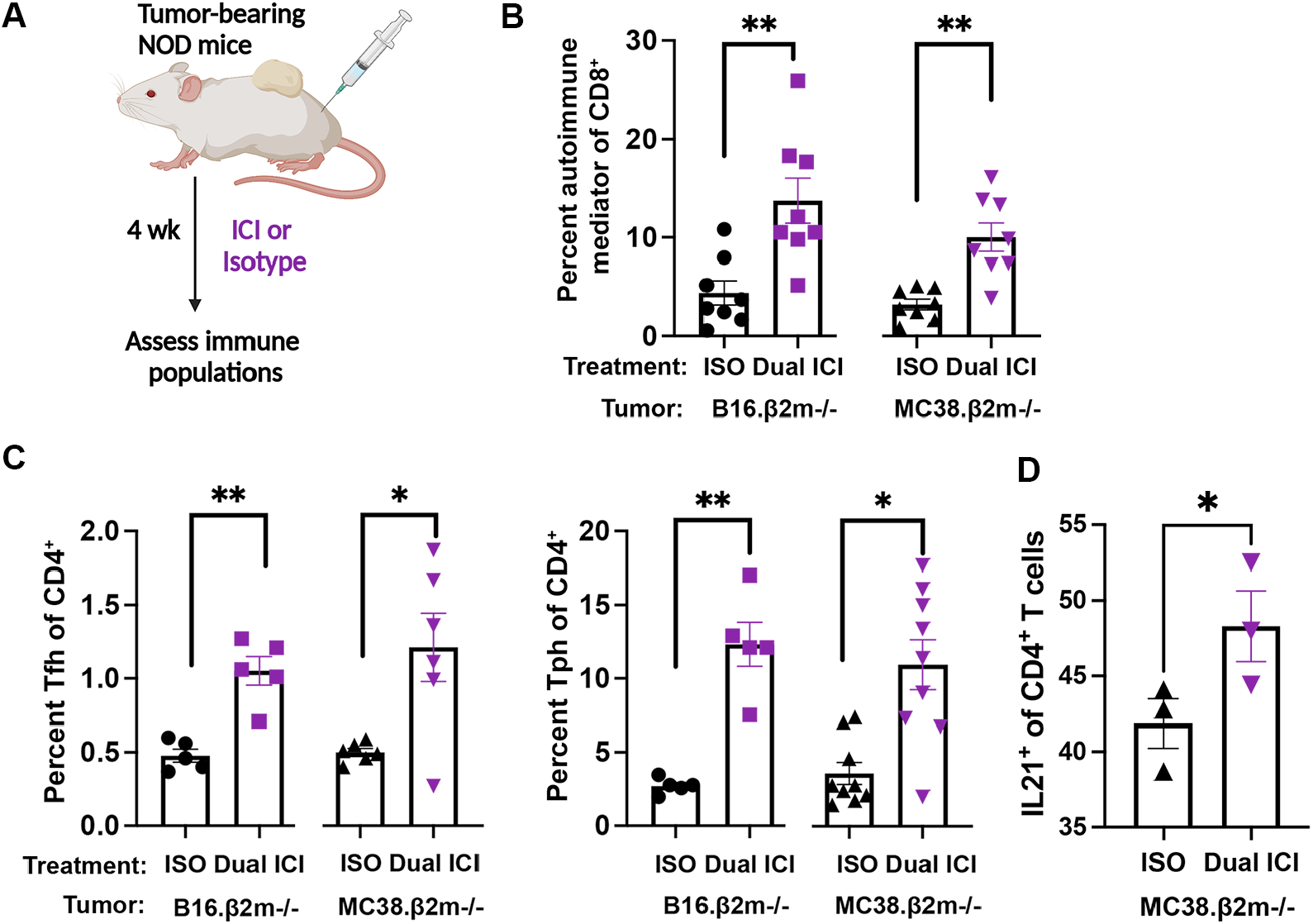
CD8^+^ Autoimmune mediator T cells and IL21^+^ Tfh and Tph cells are increased in mouse model of immune checkpoint inhibitor (ICI)-associated autoimmunity. **(A)** Schematic of mouse model, where non-obese diabetic (NOD) mice are engrafted with B16.β2M^-/-^ or MC38.β2M^-/-^ tumors and then treated with anti-PD1 + anti-CTLA4 (Dual ICI) or isotype control (ISO) for four weeks. **(B)** Frequency of autoimmune mediator (CXCR6^+^ IFNG^+^ GZMB^+^) CD8^+^ T cells in spleen. **(C)** Frequency of T follicular (Tfh: CD4^+^ PD1^+^ ICOS^+^ CXCR5^+^) and peripheral (Tph: CD4^+^ PD1^+^ ICOS^+^ CXCR5^-^) helper cells in spleen. **(D)** Frequency of IL21^+^ CD4^+^ T cells in spleen. Each point represents one animal (B-D). ANOVA with Welch correction and subsequent pairwise comparison (B,C) or Student’s t test (D), *p<0.05, **p<0.01.

### Blockade of IL21 signaling reduces CXCR6^+^ CD8^+^ autoimmune mediator T cells and protects ICI-treated mice from thyroid autoimmune infiltrates

Taken together, our data suggested a role for the IL21 Tfh/Tph – CD8^+^ autoimmune mediator axis in the development of IRAEs. Congruent with our findings in humans, recombinant IL21 increased expression of autoimmune mediator effector molecules granzyme B and IFNγ (**Fig. 6A**) and CXCR6 (**Fig. 6B**) by murine CD8^+^ T cells *in vitro*. Based upon these data, we predicted that genetic blockade of IL21 signaling in our mouse model could reduce thyroid immune infiltrates during ICI treatment (**Fig. 6C)**. We compared autoimmune infiltrates in tumor bearing (MC38.β2m^-/-^) mice with genetic deletion of the IL21 receptor (IL21RKO) to wild type (WT) controls during ICI treatment (**Fig. 6D**). Indeed, the frequency of CXCR6^+^ CD8^+^ T cells was reduced in ICI-treated IL21R KO mice compared to ICI-treated WT mice (**Fig. 6E**).

**Fig. 6.**
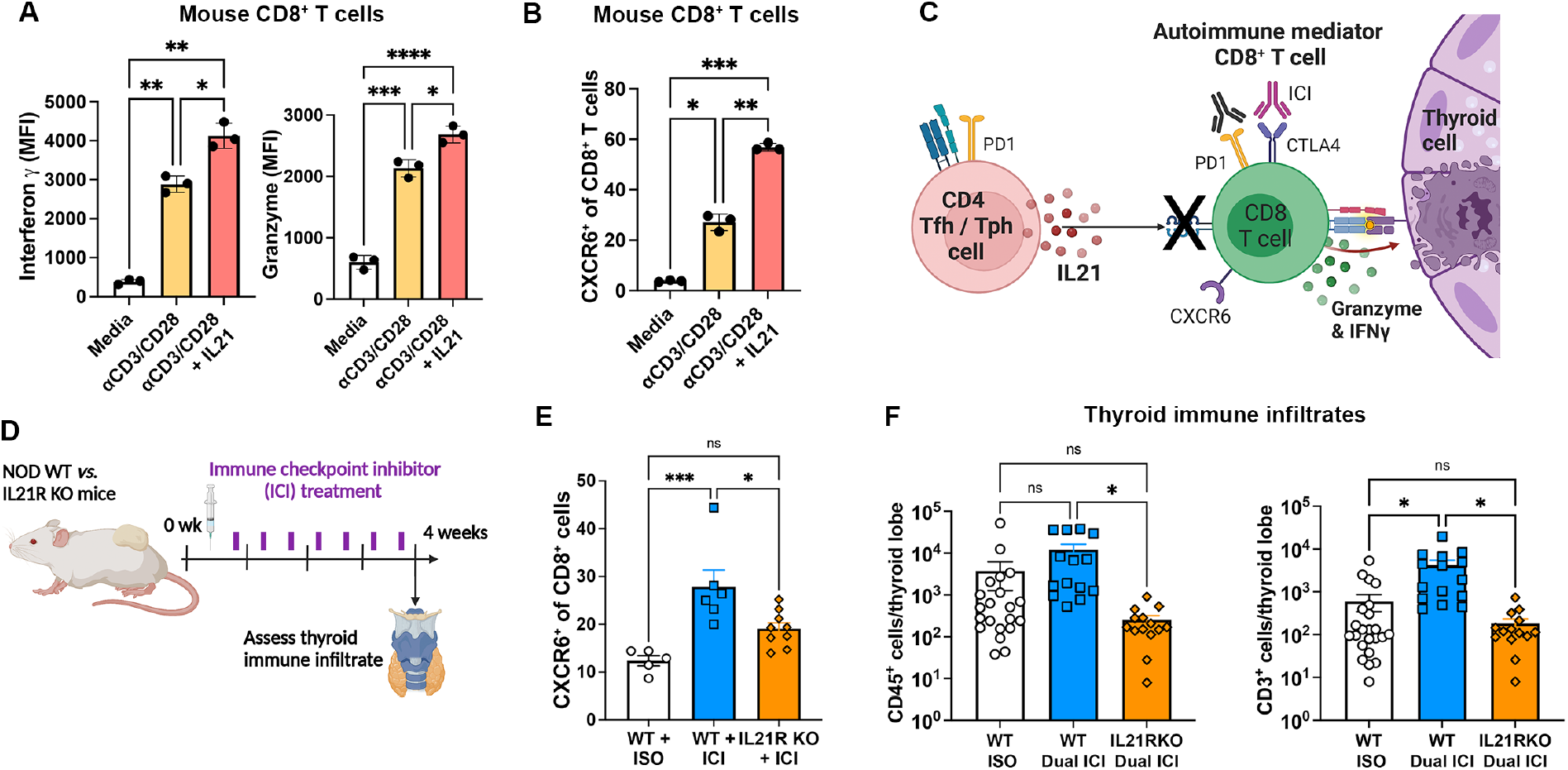
IL21 signaling is required for ICI-related thyroid autoimmunity in a mouse model. Expression of autoimmune mediator effectors interferon gamma (IFNγ) and granzyme B **(A)** and chemokine receptor CXCR6 **(B)** by murine CD8^+^ T cells stimulated 3 days *in vitro* with anti-CD3/28 and recombinant murine IL21, and then analyzed by flow cytometry. **(C)** Proposed mechanism of ICI-thyroiditis and inhibition by blockade of IL21 signaling. **(D)** Schematic of ICI treatment in NOD wild type (WT) or IL21 receptor knock out (IL21R KO) mice bearing MC38. 2m^-/-^ tumors. **(E)** Frequency of CXCR6^+^ CD8^+^ T cells in ICI-treated WT and IL21R KO spleen. **(F)** Quantification of thyroid-infiltrating CD45^+^ and CD3^+^ cells in ICI-treated mouse mice. ANOVA with Welch correction and subsequent pairwise comparison (A,B,E,F), *p<0.05, **p<0.01, ***p<0.001, ****p<0.0001.

Furthermore, IL21R KO mice were protected from ICI-associated thyroid autoimmunity compared to WT controls (**Fig. 6F**). Thyroid-infiltrating CD45^+^ and CD3^+^ immune cells were reduced after 4 weeks of anti-PD1 + anti-CTLA4 treatment in IL21R KO mice compared to ICI-treated WT mice, and similar to isotype treated WT controls (**Fig. 6F**). Thus, blockade of IL21 signaling during ICI treatment significantly reduced ICI-associated thyroid autoimmune infiltration. These data suggest that IL21-induced CD8^+^ autoimmune mediators play a crucial role in the development of ICI-thyroiditis and provide a new therapeutic target for the prevention of IRAEs in humans.

## DISCUSSION

Autoimmune toxicities remain an important limitation to the benefits and use of ICI cancer therapy. Despite significant efforts to date, strategies to reduce IRAEs have been elusive, in part due to a lack of understanding of pathogenic immune cells within the tissue microenvironment. In this study, we report for the first time the intrathyroidal immune composition of autoimmune infiltrates in human ICI-thyroiditis, one of the most common IRAEs in patients, using scRNAseq and TCR seq. Intrathyroidal infiltrates were T cell-predominant and included IL21-producing CD4^+^ Tfh and Tph populations. Comparisons to HT and healthy control samples revealed a predominant, clonally expanded CD8^+^ T cell population with an autoimmune mediator phenotype in ICI-thyroiditis. Expansion of this CD8^+^ autoimmune mediator population was recapitulated in a newly developed IRAE mouse model. Finally, our data show that CD8^+^ autoimmune mediator differentiation and thyroiditis development were dependent on IL21, since blockade of IL21R signaling prevented their differentiation and protected mice from ICI-associated thyroiditis.

CD8^+^ autoimmune mediators have recently been shown to drive pathogenic immune responses in preclinical models of autoimmune diabetes and hepatitis *(26–28)*. These cells differentiate from stem-like progenitors in lymph nodes or within the tissue to become terminal effectors with cytotoxic function. During this transition, CD8^+^ effectors down regulate TCF1 and increase expression of CXCR6, IFNγ, granzyme, PD1 and CTLA4. This autoimmune mediator population is emerging as a common effector phenotype shared across autoimmunity in different tissues. We now extend the role of CD8^+^ autoimmune mediators to ICI-thyroiditis in both humans and mice. In support of a pathogenic role in ICI, thyroiditis, thyroid-infiltrating autoimmune mediators acquire cytotoxic gene expression in conjunction with TCR clonal expansion, consistent with thyroid antigen response. Furthermore, CD8^+^ autoimmune mediators expressed PD1 and CTLA4 and therefore may be direct targets of ICI antibody binding. In contrast to HT, where T cells are kept in check by T cell PD1 binding to thyroid cell PDL1 *(43)*, in IRAEs autoreactive CD8^+^ T cells may be released from checkpoint inhibition during ICI therapy. Evidence for CD8^+^ T cell enrichment in human IRAEs was also seen in a study by Sasson et al. *(44)*, in which increased IFNγ^+^ CD8^+^ T cells were seen in patients with ICI-related colitis. Ongoing studies are interrogating these TCR sequences of clonally expanded CD8^+^ autoimmune mediators to determine antigen targets in thyroid IRAE. Clonal expansion of T cells has been linked to IRAE risk in two recent studies *(22, 45)*, including clonal expansion of CD8^+^ T cells in a mouse model of ICI-associated myocarditis *(45)*. Importantly, whether clonal expansion also occurs in human ICI-associated myocarditis is unknown as this study did not evaluate immune infiltrates in the tissues of ICI-myocarditis patients. Thus, in addition to ICI-thyroiditis, CD8^+^ autoimmune mediators may be central to IRAEs in other tissues, and more investigations at the tissue level in humans are needed.

Our data also suggest a role for IL21^+^ CD4^+^ Tfh and Tph cells in the development of ICI-thyroiditis and as a shared mechanism with spontaneous thyroid autoimmunity (*i.e*. HT). This may explain why patients with pre-existing thyroid autoantibodies are at a significantly higher risk of developing thyroid IRAEs during ICI therapy *(46)*. Wherry and colleagues recently showed increased chemokine expression (CXCL13) and B cell co-stimulation by circulating Tfh cells in anti-PD1 treated patients compared to untreated individuals *(24)*. In this study, we show that both Tfh and Tph cells infiltrate into IRAE affected thyroid tissue and propose a mechanism by which Tfh and Tph cells may contribute to autoimmunity during ICI therapy via IL21 action on CD8^+^ T cells. Specifically, recombinant IL21 drove differentiation of CD8^+^ autoimmune mediators and increased thyrotoxic activity, suggesting it as a potential target to reduce IRAEs. Indeed, we showed that genetic blockade of IL21 signaling *in vivo* was sufficient to reduce thyroid autoimmunity during ICI therapy.

We acknowledge several limitations of this study. First, the scRNAseq data presented for patients with ICI-thyroiditis encompasses human variation, including differences in primary cancer type, age and sex, and treatment history. In addition, thyroid FNA specimens yield a small number of cells from each subject and therefore confirmatory flow cytometry was not feasible for all specimens. However, these studies do provide the first in depth look into the tissue immune infiltrates of ICI-thyroiditis at greater resolution than prior flow cytometry or immunohistochemistry studies *(20, 21)*. Furthermore, *in vitro* immune assays with human cells and *in vivo* studies using our mouse model of IRAEs serve to corroborate the mechanisms inferred from our transcriptional data. Finally, to test the role of IL21 signaling in thyroid IRAE development, we used mice with genetic deletion of IL21 receptor. Future studies are needed to further elucidate the relative role of T cell, as well as B cell, subsets within this pathway.

In conclusion, we provide evidence for a diverse immune infiltrate in ICI-associated thyroid autoimmunity and a central role for CD8^+^ cytotoxic T cells. As a strategy to reduce autoimmune toxicity, we further identify a supporting role of IL21 producing CD4^+^ Tfh and Tph cells via promotion of the cytotoxic phenotype in CD8^+^ T cells. Importantly, as suggested by preclinical data in our mouse model of ICI-associated autoimmunity, blockade of IL21 signaling may provide an avenue to attenuate ICI autoimmune toxicities.

## MATERIALS AND METHODS

### Study design

Patients were prospectively enrolled from two academic medical centers (UCLA Health, USC Keck Medical Center) under IRB approved protocols (21-000633, 19-001708, HS-19-00715). Thyroid FNA specimens were collected from adult (age >18 years) patients with 1) ICI-treated cancer patients with new onset thyroid IRAE, or 2) Hashimoto’s thyroiditis. Exclusion criteria included pregnancy, history of thyroid surgery, radioactive iodine therapy, or thyroid cancer, immune modifying conditions not including solid malignancy (e.*g*. bone marrow transplantation, leukemia or lymphoma, known genetic or acquired immunodeficiency, or immune modifying medications at the time of specimen collection, excluding physiologic steroids). ICI-thyroiditis (Thyroid IRAE) was defined as new onset thyroid dysfunction (within 2 months of specimen collection) while on anti-PD1/L1 and/or anti-CTLA4 inhibitor therapy, with an overt hyperthyroid phase (low TSH and elevated FT4) followed by hypothyroidism (elevated TSH and low FT4) requiring thyroid hormone replacement. ICI-treated subjects must have received an FDA-approved ICI within the past 1 month. HT patients had hypothyroidism (elevated TSH and low FT4, and/or requirement for thyroid hormone replacement) and evidence of thyroid autoimmunity (e.*g*. thyroid autoantibody presence, anti-TPO or anti-Tg); imaging findings if available were consistent with HT. Patients may have had a cytologically-proven benign thyroid nodule. Patient demographic, treatment and clinical immune data are summarized in **Table S1**. Thyroid FNA specimens were collected into RPMI media under ultrasound guidance using 4 passes with a 25 or 27 gauge needle by M.G.L, T.E.A., or P.F.

### Single cell RNA sequencing

For human scRNAseq of thyroid FNA specimens, single cell suspensions were stained with fluorescence-conjugated antibodies to CD45 and viability dye DAPI. Single live CD45^+^ immune cells were collected and sorted, then submitted for 10x 5’ and TCR sequencing. Cell preparation, library preparation, and sequencing were carried out according to Chromium product-based manufacturer protocols (10X Genomics). Sequencing was carried out on a Novaseq6000 S2 2×50bp flow cell (Illumina) utilizing the Chromium single-cell 5′ and TCR gene expression library preparation (10X Genomics), per manufacturer’s protocol at the Cedars Sinai Medical Center Genomics Core.

### Data processing of scRNAseq libraries

After sequencing, the scRNAseq reads were demultiplexed and aligned with Cell Ranger versionv3.0.0 or higher (10X Genomics) to the GRCh38 human reference genome and quantified using the 10x Genomics CellRanger count software (10x Genomics). Filtered output matrices that contained only barcodes with unique molecular identifier (UMI) counts that passed the threshold for cell detection were used for downstream analysis.

### Normalization, Principal Component analysis, and UMAP Clustering

Downstream analysis was done with Seurat v3 or higher *(47)*. Only cells with a mitochondrial gene percentage less than 30% and 200 features were included in downstream analysis. Scores for S and G2/M cell cycle phases were assigned using the Seurat CellCycleScoring function following the standard Seurat pipeline *(48)*. UMI counts were log normalized, and the top 2000 variable genes were determined using the variance-stabilizing transformation (vst) method. All genes were scaled and centered using the ScaleData function, and principal component analysis (PCA) was run for the data using the predetermined variable genes. To group cells into clusters, a K-nearest neighbors graph function (implemented in the Seurat package), followed by a modularity-optimizing function using the Louvain algorithm was used. For clustering, 30 PC dimensions were included and the resolution parameter was set to 0.8. Cell-type clusters were visualized using uniform manifold approximation and projection (UMAP) to reduce dimensionality and allow for the cells to be visualized on a 2-D plot. Subclustering of CD8 T cells was done by selection of cells with *CD8A* and *CD8B* expression >1 and *CD4, NCR1*, and *TRDC* expression <0.5, and then reclustered with 10 PC dimensions and a resolution of 0.4. CD4 T cells were subclustered by gene expression *CD4*>1 and *CD8A* < 0.5 and *TRDC* <0.5 and then reclustered with 10 PC dimensions and a resolution of 0.6.

### Differential Expression and Marker Gene Identification

The Seurat FindMarkers function was used to generate the top upregulated genes for each cluster using a Wilcoxon Rank Sum Test to identify differentially expressed genes across clusters. Marker genes were filtered by a minimum of detectable expression in 25% of the cells in the target group and minimum log2 fold change of 0.25. The markers generated by these functions were compared to markers for known cell types to assign identities to the different clusters. Specifically, variably expressed gene sets for each cluster were queried in curated, publicly available databases for putative cell population identity: Enrichr *(49)* and the Immunological Genome Project database *(50)*. Differentially expressed genes across conditions were identified using the same function and parameters. Cells from proliferating clusters, stressed/dying clusters, doublets, and unknown clusters were excluded from this analysis. Significant DEG between conditions, defined as log fold change > 0.5 and P value < 0.05, were used for gene-set enrichment analysis using Metascape *(51)*.

### Trajectory analysis

Trajectory inference was performed on the subsetted CD4 and CD8 T-cells using Slingshot*(52)*, which uses a cluster-based MST to identify global lineage structure, then fits smooth curves to lineages to infer pseudotime values for each cell. Lineages were determined using the UMAP reduction calculated using Seurat. Starting clusters for each lineage were pre-selected for naïve T cells. To test for significant differences in pseudotime distributions between conditions, asymptotic Kolmogorov-Smirnov tests were conducted between pairwise comparisons across the three conditions (HT, IRAE, and Healthy Control) for each lineage.

### TCR clonotype analysis and mapping

To identify and further analyze TCR information produced by 10x Genomics Cell Ranger pipeline, scRepertoire *(53)* was used to assign clonotypes and discern TCR dynamics within the data. Moreover, by applying the frequency for each clonotype sequence, the top clonotypes of each condition were mapped onto the UMAP of single-cell expression datasets from Seurat to visualize the TCR information on a cluster level. The clonal expansion data was also integrated with the single-cell expression datasets from Seurat to compare gene expression between clonally expanded T cells and non-clonally expanded T cells. Differences in clonal expansion and clone overlap between clusters was determined using STARTRAC *(54)*.

### Cell lines and media

Cell lines used in these studies included NThyOri-3.1, a human thyroid follicular cell line generated by SV40 transformation of normal thyroid cells, obtained from the American Type Culture Collection, and murine colon tumor model MC38 with genetic deletion of beta 2 microglobulin (MC38.β2M^-/-^) and melanoma tumor model (B16.β2M^-/-^), kindly gifted from Dr. Antoni Ribas *(55)*. Tumor cell line authenticity was performed by surface marker analysis performed at ATCC or in our laboratory and phenotype was regularly monitored in culture. Cell lines and primary immune cells and cell lines were cultured in RPMI complete media [supplemented with 10% fetal bovine serum (FBS), 2mM L-glutamine, 1mM HEPES, non-essential amino acids, and antibiotics (penicillin and streptomycin)], with 50uM beta-mercaptoethanol (2ME).

### *In vitro* assessment of primary human immune cells

Peripheral blood mononuclear cells (PBMC) were isolated from blood by density gradient centrifugation (Ficoll-Hypaque). CD8^+^ T cells were separated by magnetic bead separation (Miltenyi Biotec). For T cell stimulation assays, PBMC or isolated T cells were cultured at 5×10^5^ cells/well in 12 well plates in complete media with 2ME. Stimulation was provided by human anti-CD3/CD28 Dynabeads (Invitrogen) and recombinant human IL21 at 100ng/mL (PeproTech) or vehicle control, as indicated. Isolated CD8 T cells were additionally cultured in the presence of recombinant IL-2 at 50 units/mL. Cytokines were refreshed at day 3 and cells analyzed by flow cytometry on day 5. Experiments were repeated at least twice.

To test the effect of T cell supernatants on thyroid cell viability, NThy-Ori 3.1 cells were cultured with day 3 supernatant from human CD8^+^ T cell *in vitro* cultures diluted 1:1 with fresh complete media. Cell viability was evaluated by staining with DAPI and analysis by flow cytometry at 24hr. Experiments repeated at least twice.

### Mouse studies

Animal studies were approved by the University of California Los Angeles Animal Research Committee (Protocol C21-039). NOD/ShiLtJ (NOD) and NOD/Il21r-KO (IL21R KO) mice were obtained from the Jackson Laboratory. Male and female mice were used in equal proportions. Mice were used at 4-6-weeks of age unless otherwise noted. Mice were housed in a specific pathogen-free barrier facility at the University of California Los Angeles. Mice in different experimental groups were co-housed.

### Immune checkpoint inhibitor treatment of mice

Mice were inoculated subcutaneously with 6×10^5^ MC38.β2M^-/-^ or B16.β2M^-/-^ tumor cells in the right flank and randomized into treatment groups when tumor volumes reached 50-80mm^3^. Mice were randomized to twice weekly treatment with anti-mouse CTLA4 (clone 9D9) and PD1 (clone RPM1-14) or isotype control antibodies (2A3, MPC-11), at 10 mg/kg/dose intraperitoneally *(i.p.)* for four weeks, as described previously*(9)*. ICI were from BioXcell. During treatment mice were monitored daily for activity and appearance, and twice weekly for weight and glucosuria. Mice developing glucosuria were treated with 10 units of subcutaneous NPH insulin daily. After four weeks of ICI treatment, mice were euthanized, blood collected by retro-orbital bleed, then perfused with 10mL of phosphate buffered saline (PBS). For evaluation of immune infiltrates by flow, fresh thyroid glands were dissected away from surrounding trachea and lymphoid tissue, digested in collagenase type IV (1mg/mL in 2% FBS in PBS) at 37°C for 20 minutes, then mechanically dissociated by passage through a 40um filter. Spleen cells were isolated by mechanical dissociation and passage through a 40um filter. Predetermined endpoints for euthanasia before four weeks included >20% weight loss and glucosuria not resolved by insulin therapy, as per IACUC protocols. Immediately after euthanasia and perfusion with sterile PBS, fresh tissues were dissociated for analysis of immune infiltrates by flow cytometry.

### *In vitro* assessment of primary murine immune cells

Splenocytes were isolated from healthy NOD.WT mice by mechanical dissociation, CD8^+^ T cells isolated, and cultured at 5×10^5^ cells/well in 12 well plates in complete media with 2ME. Stimulation was provided by mouse anti-CD3/CD28 Dynabeads (Invitrogen) and recombinant murine IL21 at 100ng/mL (PeproTech) or vehicle control, as indicated. Cells were evaluated on day 3. Experiments were repeated at least twice.

### Flow Cytometry

For staining, single cell suspensions were resuspended in FACS buffer (0.5mM EDTA, 2% FBS in PBS) at 10^6^cells/mL and stained with fluorescence conjugated antibodies. For intracellular staining, after surface staining, cells were fixed and permeabilized using cytoplasmic fixation and permeabilization kit (BD), per manufacturer instructions, with 20 min fixation at 4°C. To assess intracellular cytokines, cells were incubated in complete RPMI media with 50uM 2ME for 4 hours with ionomycin (1ug/mL) and PMA (50ng/mL) in the presence of Brefeldin A prior to staining. Viability dye DAPI was added prior to analysis where indicated. Cells were then washed twice in FACS buffer and analyzed by flow cytometry on an Attune NxT 6 cytometer (ThermoFisher). Antibodies used are shown in **Supplemental Methods**. Gating strategies are shown in **Fig. S5**. Cell counts are shown as relative frequency of live, gated single cells unless otherwise noted. For determination of infiltrating cells per thyroid lobe, absolute cell counts per thyroid lobe were determined by a calculation of cell count x fraction of thyroid analyzed to estimate a total cell count per thyroid lobe, as done previously *(9)*.

### Statistical analysis

Analyses of scRNAseq data were performed in RStudio (v2021.09.2) as described above. Statistical analyses for human and mouse studies were performed with GraphPad Prism software (v9.3.1). Comparisons among multiple groups for continuous data were made using ANOVA or ANOVA with Welch correction with no assumption for equal variances, with subsequent pairwise comparisons. Comparisons between two groups were done by Student’s t test. When multiple comparisons were performed, adjusted p-values are shown. Significance was defined as α = 0.05.

## List of Supplementary Materials

### Supplemental Materials and Methods

**Fig. S1 to S5**

**Table S1**. Clinical characteristics of patients.

**Data file S1**. Differential gene expression among cell clusters for scRNAseq data.

## Author contributions

Conceptualization: M.G.L., A.D., A.R., T.E.A., M.A.S.

Methodology: M.G.L., W.H., A.R., T.E.A., M.A.S.

Investigation: M.G.L., Y.Z., A.I.H., N.H., J.O., L.N.S., H.C., A.Y.P., R.Y.T., K.K., P.Y.

Visualization: M.G.L., Y.Z., H.C., A.Y.P.

Funding acquisition: M.G.L., T.E.A., M.A.S.

Project administration: M.G.L., M.A.S.

Supervision: M.G.L., M.A.S.

Writing – original draft: M.G.L., Y.Z., N.H., L.N.S., A.Y.P.

Writing – review & editing: J.O., W.H., H.C., A.D., T.E.A., P.F., A.R., M.A.S.

## Data and materials availability

Data associated with figures are available from the corresponding author upon reasonable request. The datasets for single cell sequencing generated during and analyzed during the current study are available in the Gene Expression Omnibus (GEO) repository under accession number GSE218743 (https://www.ncbi.nlm.nih.gov/geo/).

## Figures

**Table S1.**
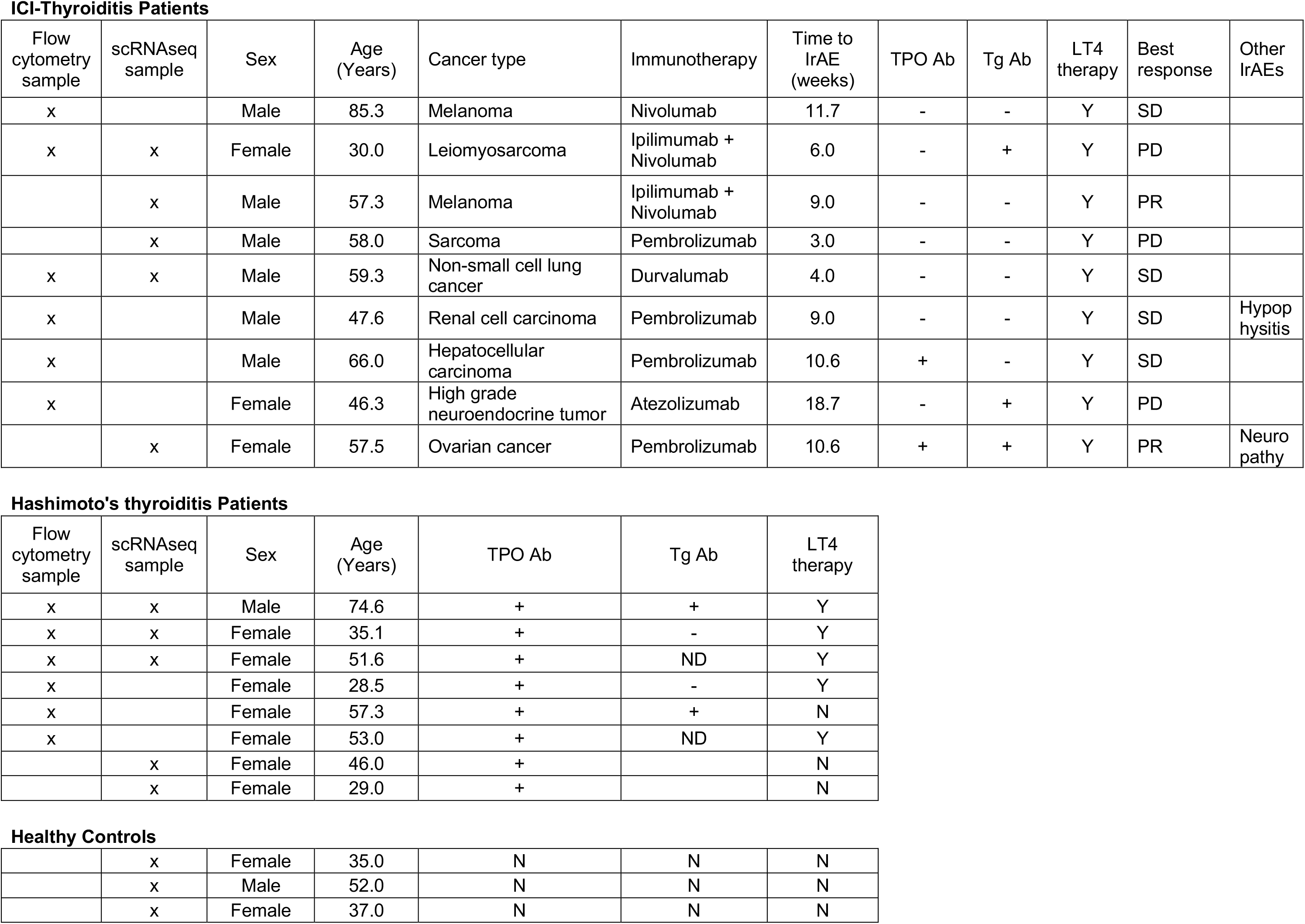
Clinical characteristics of patients. Individuals with immune checkpoint inhibitor (ICI)-associated thyroiditis, Hashimoto’s thyroiditis (HT), or no thyroid disease were enrolled for thyroid fine needle aspiration (FNA) specimen collection. Demographic, diagnosis and treatment history, thyroid autoantibody [thyroid peroxidase antibody (TPO Ab) or thyroglobulin antibody (Tg Ab)], thyroid function, and thyroid hormone replacement status data are shown. Tumor response shown as stable disease (SD), partial response (PR), or progression of disease (PD).

## Supplemental Materials and Methods

Antibodies used in flow cytometry experiments

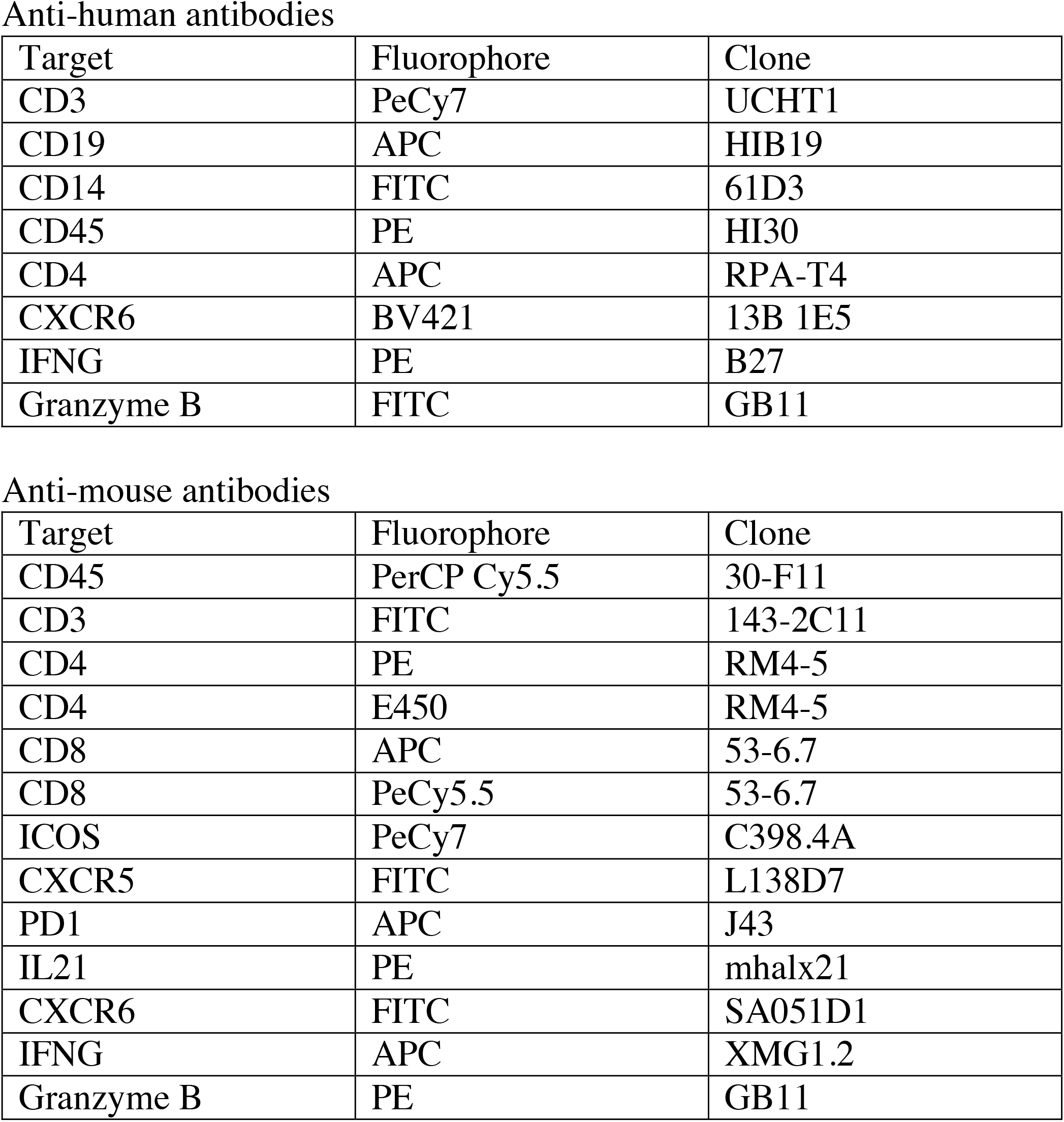

## Supplemental Figures

**Fig. S1.**
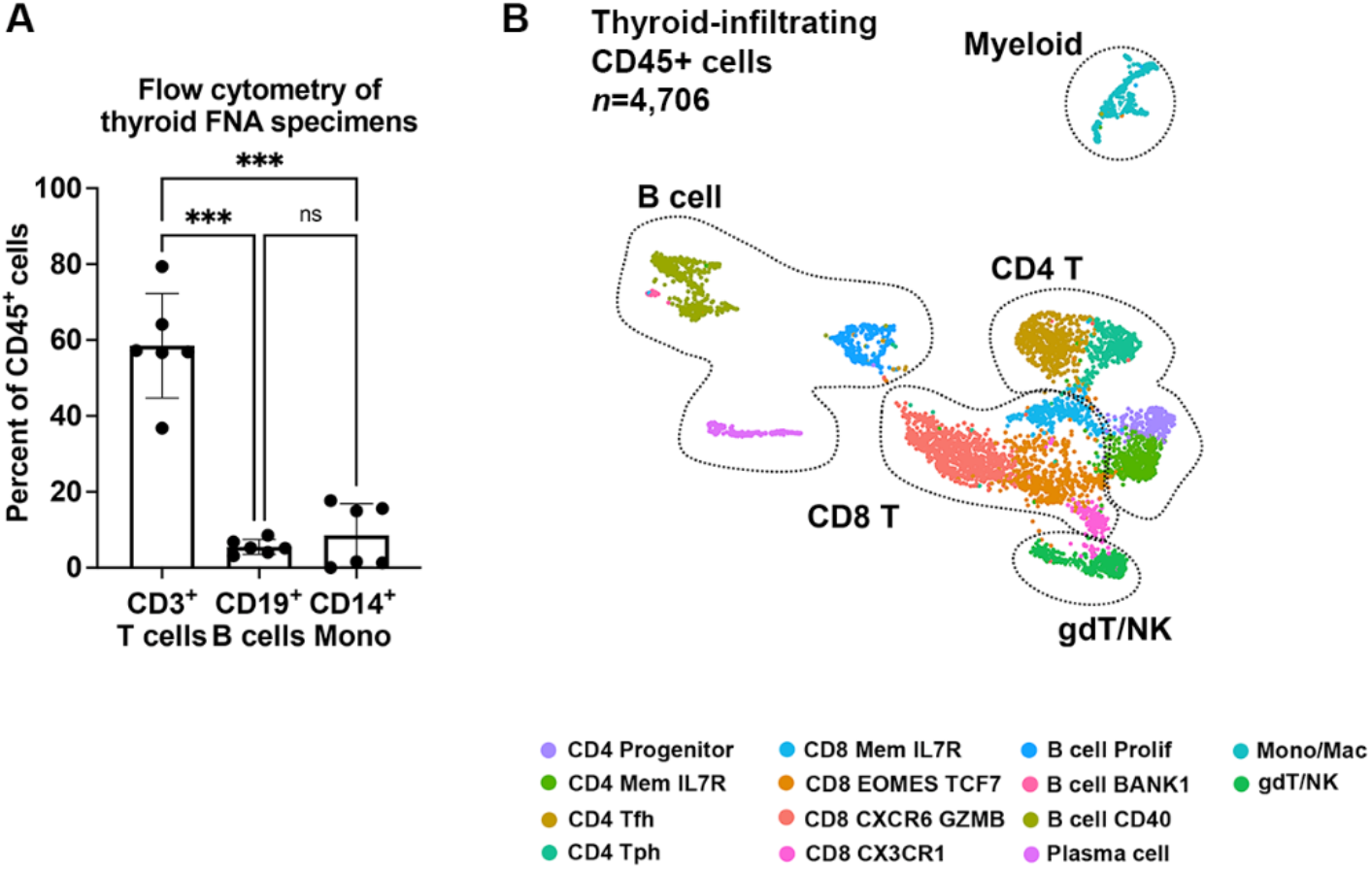
Immune composition of ICI-thyroiditis. **(A)** Immune populations by flow cytometry in thyroid FNA specimens (*n*=6 patients). ANOVA with Welch correction, ***p<0.001. **(B)** Single cell RNA sequencing analysis of thyroid infiltrating CD45^+^ immune cells (*n*=5 patients), UMAP.

**Fig. S2.**
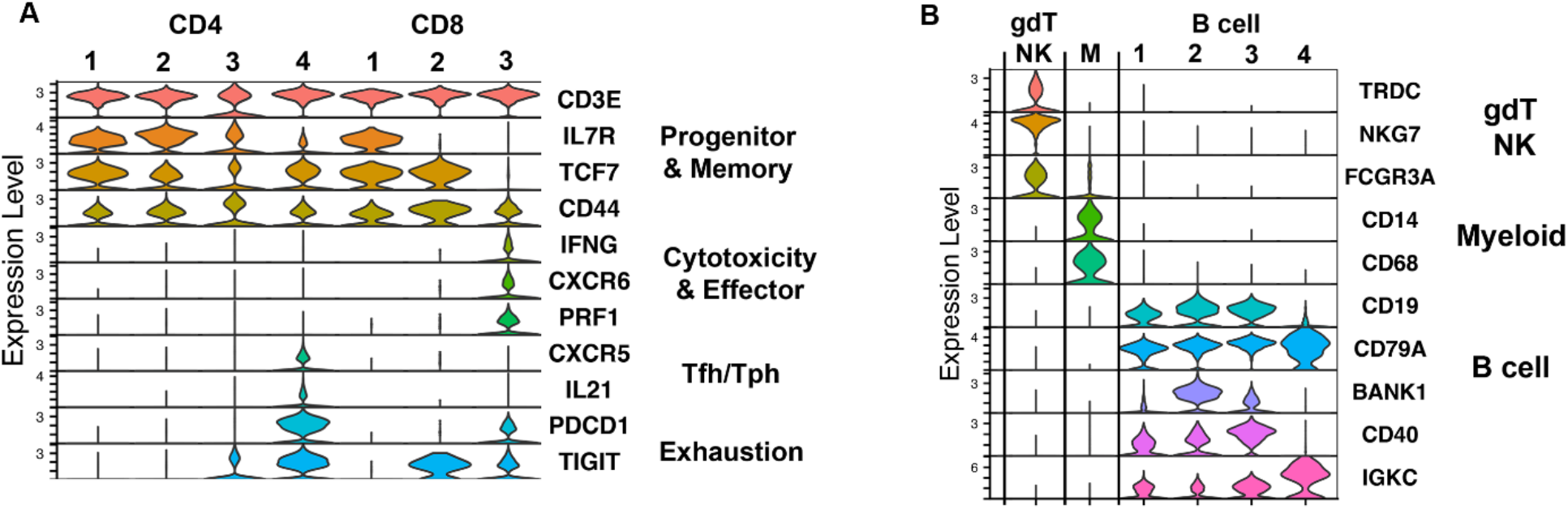
Gene expression across immune cell clusters in ICI-thyroiditis. Stacked violin plot showing gene expression by cell cluster for CD4 and CD8 T cells **(A)** and other immune cell populations **(B)**, including gamma delta T (gdT), natural killer (NK), myeloid (M), and B cell populations.

**Fig. S3.**
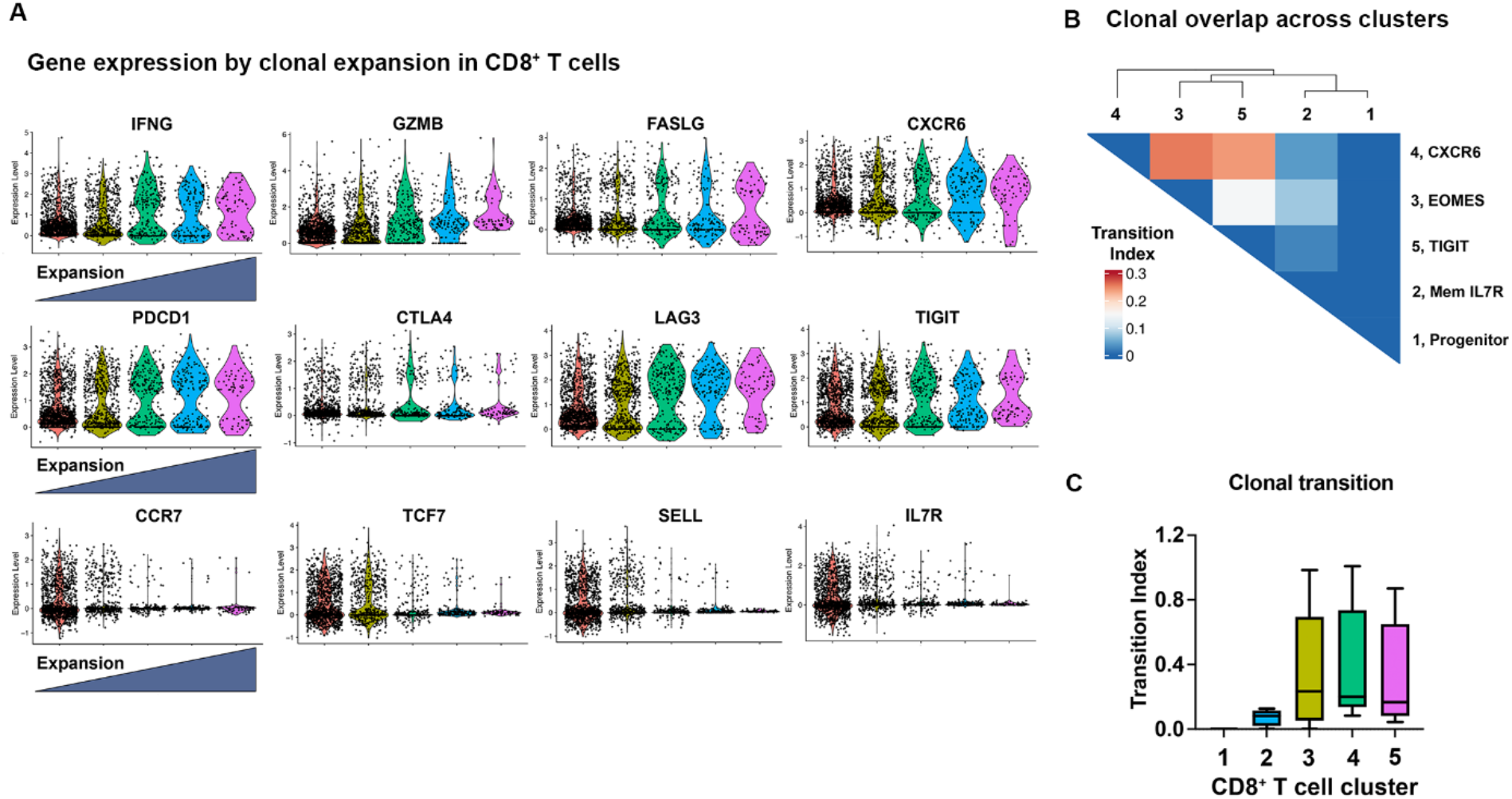
Clonally expanded CD8^+^ T cells in ICI-thyroiditis. **(A)** Violin plots showing gene expression by extent of clonal expansion. **(B)** Clonal overlap and **(C)** clonal transition index across CD8^+^ T cell clusters by STARTRAC. ANOVA, p=ns.

**Fig. S4.**
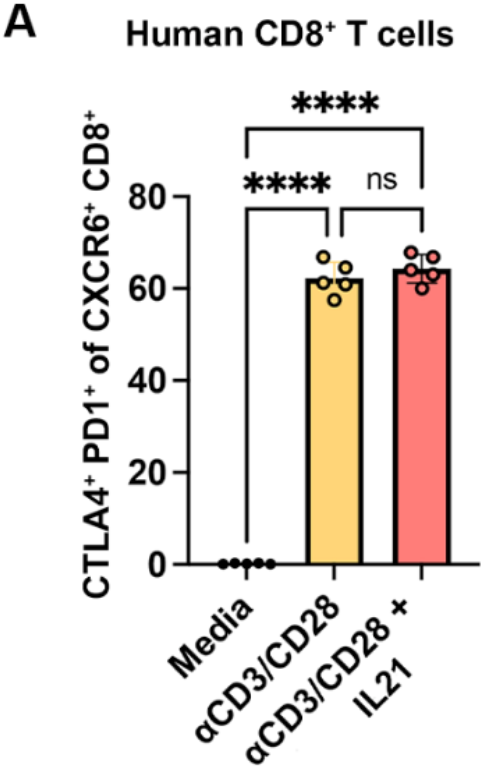
IL21-induced human CXCR6^+^ CD8^+^ T cells express checkpoint proteins. **(A)** Expression of programmed death protein (PD1) and cytotoxic T lymphocyte antigen (CTLA) 4 on *in vitro* anti-CD3/28 + IL21-stimulated human CXCR6^+^ CD8^+^ T cells by flow cytometry. ANOVA with Welch correction and subsequent pairwise comparisons. ****p<0.0001.

**Fig. S5.**
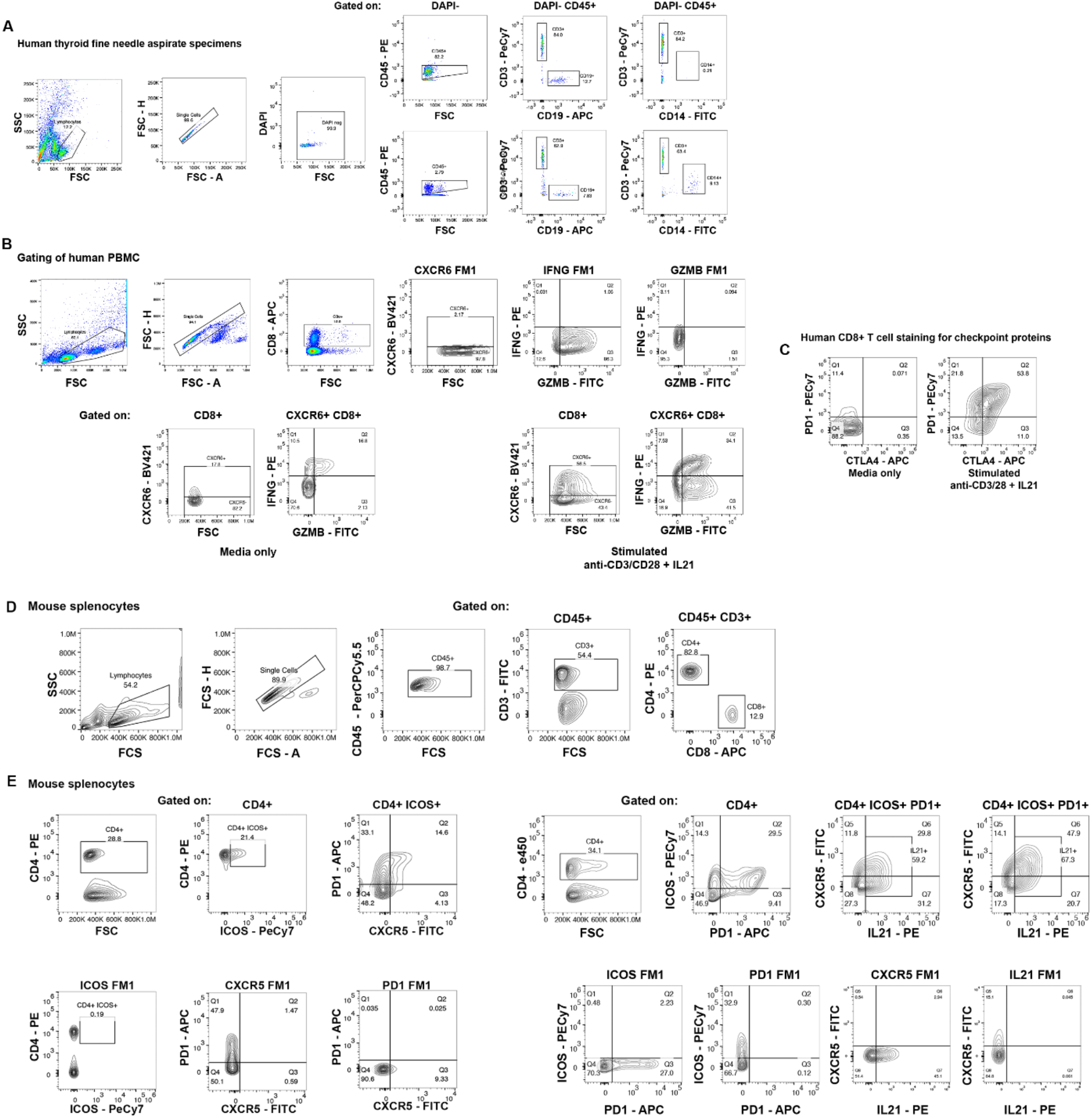
Gating strategy for flow cytometry. Representative flow cytometry plots and gating strategies for human **(A-C)** and mouse **(D-E)** immune cells.

