## Supplemental Materials for "Clonally-Expanded, Thyrotoxic Autoimmune Mediator CD8^+^ T cells Driven by IL21 Contribute to Checkpoint Inhibitor Thyroiditis"

**ICI-Thyroiditis Patients**

| Flow cytometry sample | scRNAseq sample | Sex | Age (Years) | Cancer type | Immunotherapy | Time to IrAE (weeks) | TPO Ab | Tg Ab | LT4 therapy | Best response | Other IrAEs |
|---|---|---|---|---|---|---|---|---|---|---|---|
| x | | Male | 85.3 | Melanoma | Nivolumab | 11.7 | - | - | Y | SD | |
| x | x | Female | 30.0 | Leiomyosarcoma | Ipilimumab + Nivolumab | 6.0 | - | + | Y | PD | |
| | x | Male | 57.3 | Melanoma | Ipilimumab + Nivolumab | 9.0 | - | - | Y | PR | |
| | x | Male | 58.0 | Sarcoma | Pembrolizumab | 3.0 | - | - | Y | PD | |
| x | x | Male | 59.3 | Non-small cell lung cancer | Durvalumab | 4.0 | - | - | Y | SD | |
| x | | Male | 47.6 | Renal cell carcinoma | Pembrolizumab | 9.0 | - | - | Y | SD | Hypop hysitis |
| x | | Male | 66.0 | Hepatocellular carcinoma | Pembrolizumab | 10.6 | + | - | Y | SD | |
| x | | Female | 46.3 | High grade neuroendocrine tumor | Atezolizumab | 18.7 | - | + | Y | PD | |
| | x | Female | 57.5 | Ovarian cancer | Pembrolizumab | 10.6 | + | + | Y | PR | Neuro pathy |

**Hashimoto's thyroiditis Patients**

| Flow cytometry sample | scRNAseq sample | Sex | Age (Years) | TPO Ab | Tg Ab | LT4 therapy |
|---|---|---|---|---|---|---|
| x | x | Male | 74.6 | + | + | Y |
| x | x | Female | 35.1 | + | - | Y |
| x | x | Female | 51.6 | + | ND | Y |
| x | | Female | 28.5 | + | - | Y |
| x | | Female | 57.3 | + | + | N |
| x | | Female | 53.0 | + | ND | Y |
| | x | Female | 46.0 | + | | N |
| | x | Female | 29.0 | + | | N |

**Healthy Controls**

| | | | | | | |
|---|---|---|---|---|---|---|
| | x | Female | 35.0 | N | N | N |
| | x | Male | 52.0 | N | N | N |
| | x | Female | 37.0 | N | N | N |

**Table S1. Clinical characteristics of patients.** Individuals with immune checkpoint inhibitor (ICI)-associated thyroiditis, Hashimoto's thyroiditis (HT), or no thyroid disease were enrolled for thyroid fine needle aspiration (FNA) specimen collection. Demographic, diagnosis and treatment history, thyroid autoantibody [thyroid peroxidase antibody (TPO Ab) or thyroglobulin antibody (Tg Ab)], thyroid function, and thyroid hormone replacement status data are shown. Tumor response shown as stable disease (SD), partial response (PR), or progression of disease (PD).

**Supplemental Materials and Methods**

Antibodies used in flow cytometry experiments

Anti-human antibodies

| Target | Fluorophore | Clone |
|---|---|---|
| CD3 | PeCy7 | UCHT1 |
| CD19 | APC | HIB19 |
| CD14 | FITC | 61D3 |
| CD45 | PE | HI30 |
| CD4 | APC | RPA-T4 |
| CXCR6 | BV421 | 13B 1E5 |
| IFNG | PE | B27 |
| Granzyme B | FITC | GB11 |

Anti-mouse antibodies

| Target | Fluorophore | Clone |
|---|---|---|
| CD45 | PerCP Cy5.5 | 30-F11 |
| CD3 | FITC | 143-2C11 |
| CD4 | PE | RM4-5 |
| CD4 | E450 | RM4-5 |
| CD8 | APC | 53-6.7 |
| CD8 | PeCy5.5 | 53-6.7 |
| ICOS | PeCy7 | C398.4A |
| CXCR5 | FITC | L138D7 |
| PD1 | APC | J43 |
| IL21 | PE | mhalx21 |
| CXCR6 | FITC | SA051D1 |
| IFNG | APC | XMG1.2 |
| Granzyme B | PE | GB11 |

**Supplemental Figures**

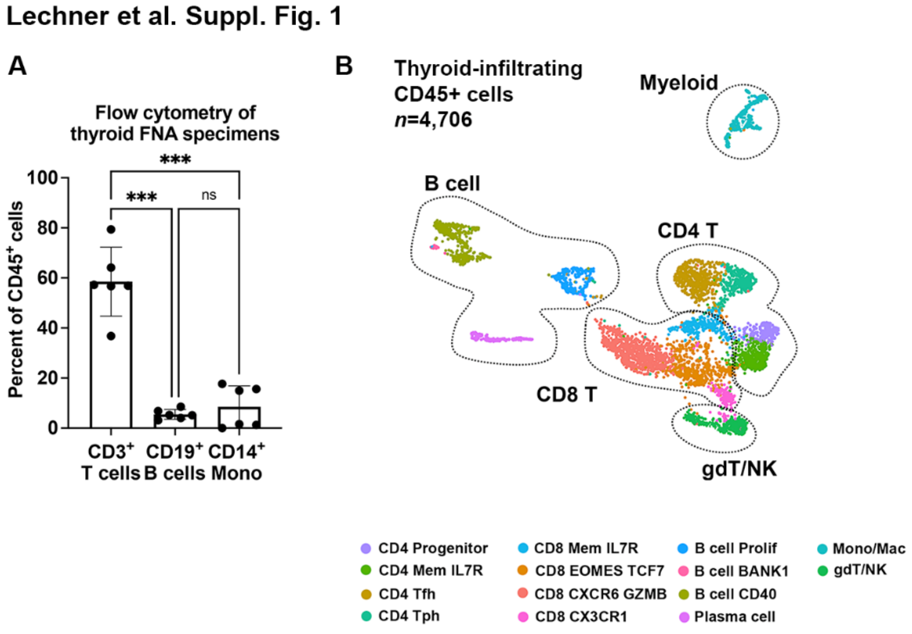


**Fig S1. Immune composition of ICI-thyroiditis (A)** Immune populations by flow cytometry in thyroid FNA specimens (*n*=6 patients). ANOVA with Welch correction, ***p<0.001. **(B)** Single cell RNA sequencing analysis of thyroid infiltrating $CD45^+$ immune cells (*n*=5 patients), UMAP.

Lechner et al. Suppl. Fig 2.

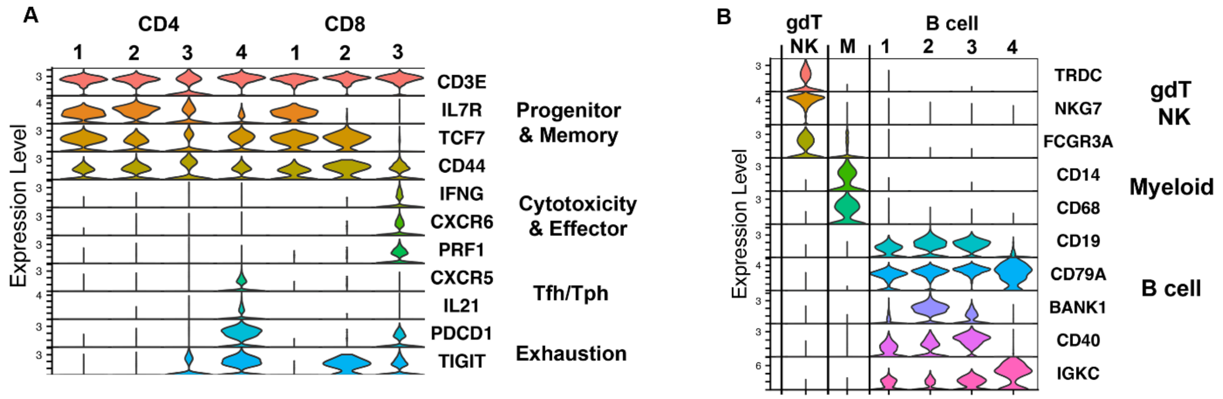


**Fig S2. Gene expression across immune cell clusters in ICI-thyroiditis.** Stacked violin plot showing gene expression by cell cluster for CD4 and CD8 T cells **(A)** and other immune cell populations **(B)**, including gamma delta T (gdT), natural killer (NK), myeloid (M), and B cell populations.

Lechner et al. Suppl. Fig. 3

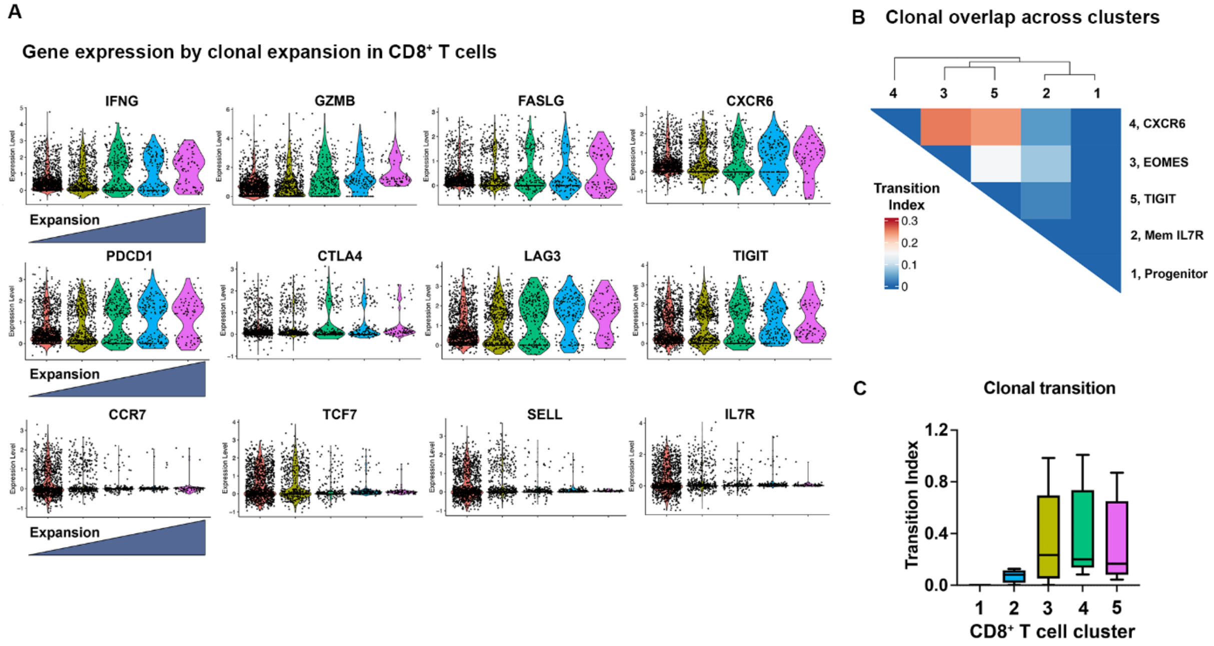


**Fig. S3**. **Clonally expanded CD8⁺ T cells in ICI-thyroiditis. (A)** Violin plots showing gene expression by extent of clonal expansion. **(B)** Clonal overlap and **(C)** clonal transition index across CD8⁺ T cell clusters by STARTRAC. ANOVA, p=ns.

Lechner et al. Suppl. Fig. 4

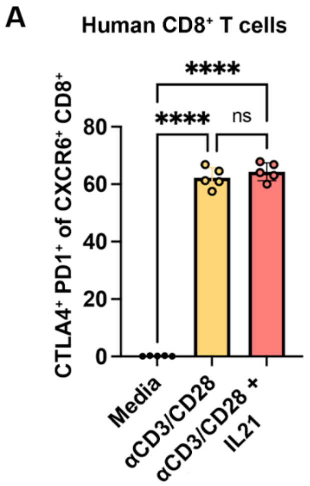


**Fig. S4**. **IL21-induced human CXCR6$^+$ CD8$^+$ T cells express checkpoint proteins. (A)** Expression of programmed death protein (PD1) and cytotoxic T lymphocyte antigen (CTLA) 4 on *in vitro* anti-CD3/28 + IL21-stimulated human CXCR6$^+$ CD8$^+$ T cells by flow cytometry. ANOVA with Welch correction and subsequent pairwise comparisons. ****$p<0.0001$.

Lechner et al. Suppl. Fig. 5

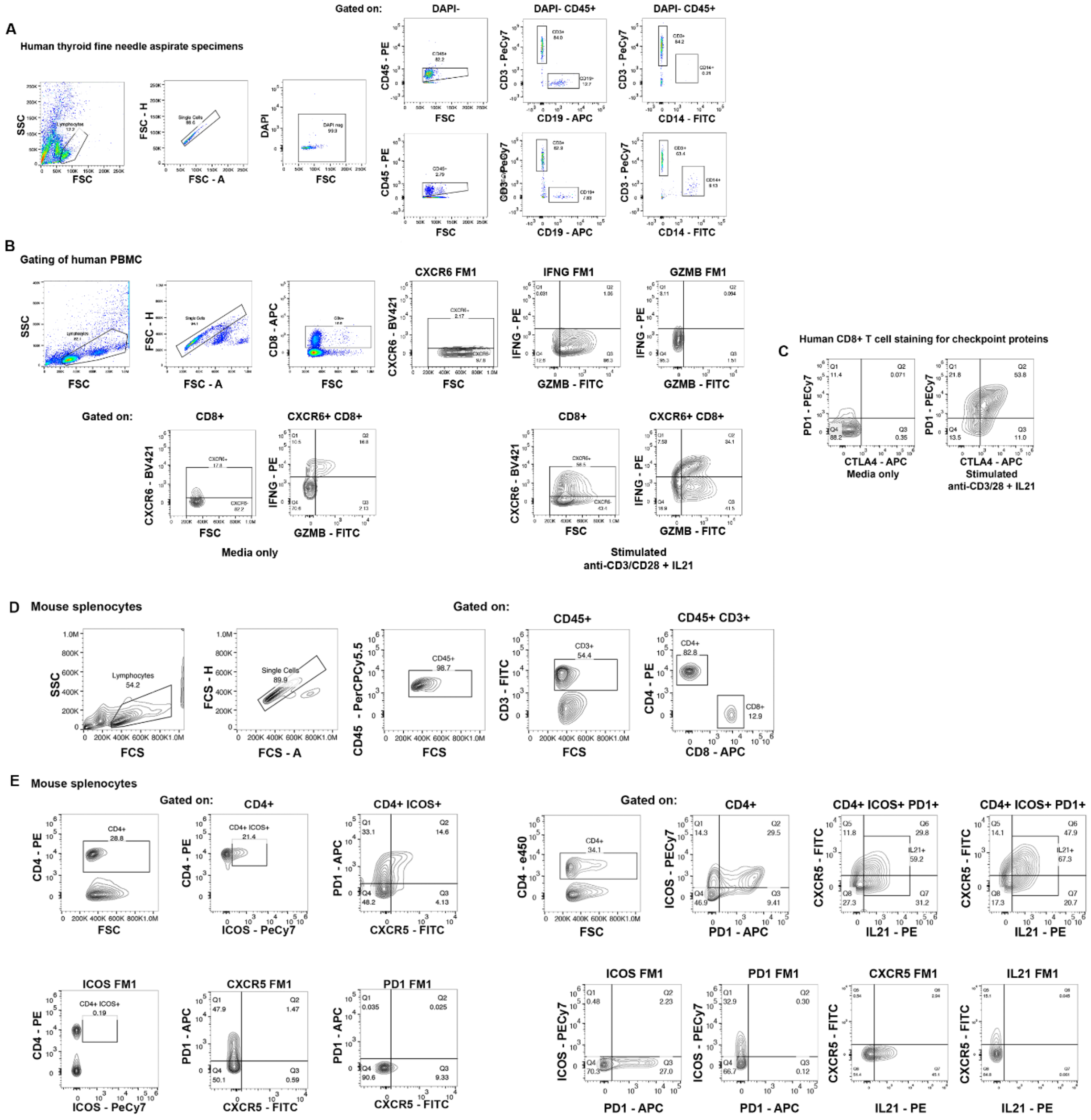

**Fig. S5**. Gating strategy for flow cytometry. Representative flow cytometry plots and gating strategies for human **(A-C)** and mouse **(D-E)** immune cells.
